# Harnessing PROTAC technology to combat stress hormone receptor activation

**DOI:** 10.1101/2023.03.17.533120

**Authors:** Mahshid Gazorpak, Karina M. Hugentobler, Dominique Paul, Pierre-Luc Germain, Kei Matthis, Remo Rudolf, Sergio Mompart Barrenechea, Miriam Kretschmer, Vincent Fischer, Xiaohan Xue, Mattia Privitera, Iryna Ivanova, Andreas Hierlemann, Onno C. Meijer, Erick M. Carreira, Johannes Bohacek, Katharina Gapp

**Affiliations:** Laboratory of Epigenetics and Neuroendocrinology, Institute for Neuroscience, Department of Health Science and Technology, ETH Zurich, 8057 Zurich, Switzerland; Neuroscience Center Zurich, ETH Zurich and University of Zurich, Switzerland; Laboratory of Organic Chemistry, Department of Chemistry and Applied Biosciences, ETH Zurich, Zurich, Switzerland; Lab of Statistical Bioinformatics, University of Zürich, Switzerland; Computational Neurogenomics, Institute for Neuroscience, Department of Health Science and Technology, ETH Zurich, 8057 Zurich, Switzerland; Bio Engineering Laboratory, Institute for Department of Biosystems Science and Engineering, ETH Zurich, 4058 Basel, Switzerland; Laboratory of Molecular and Behavioral Neuroscience, Institute for Neuroscience, Department of Health Science and Technology, ETH Zurich, 8057 Zurich, Switzerland; Department of Medicine, Division of Endocrinology, Leiden University Medical Center, Leiden, the Netherlands

**Keywords:** Glucocorticoid receptor, proteolysis targeting chimeras (PROTACs), targeted protein degradation (TPD), stress response, HPA axis, neuropsychiatric disorders, depression, anxiety, dexamethasone, mifepristone

## Abstract

Counteracting the overactivation of glucocorticoid receptors (GR) is an important therapeutic goal in stress-related psychiatry and beyond. The only clinically approved GR antagonist lacks selectivity and induces unwanted side effects. To complement existing tools of small-molecule-based inhibitors, we present a highly potent, novel catalytically-driven GR degrader, KH-103, based on proteolysis-targeting chimera technology. This selective degrader enables immediate and reversible GR depletion that is independent of genetic manipulation and circumvents transcriptional adaptations to inhibition. KH-103 achieves passive inhibition, preventing agonistic induction of gene expression, and significantly averts the GR’s genomic effects compared to two currently available inhibitors. Application in primary-neuron cultures revealed the dependency of a glucocorticoid-induced increase in spontaneous calcium activity on GR. Finally, we present a proof of concept for application in-vivo. KH-103 opens opportunities for a more lucid interpretation of GR functions with translational potential.

## Introduction

Excessive signaling by glucocorticoid (GC) hormones is linked to many pathologies^1^. Exposure to stressful situations triggers activation of the hypothalamus-pituitary-adrenal (HPA) axis, which results in the release of GCs into the bloodstream. Under healthy conditions, GCs support adaptation through modulation of all major organ systems, including the brain, in large measure via the glucocorticoid receptor (GR)^2,3^. GRs act as a transcription factor (TF): upon activation, cytosolic GR translocates to the nucleus and binds directly or indirectly to genomic response elements as a homo or heterodimer^4^. This process can result in the promotion or inhibition of transcription and/or alter chromatin accessibility to prime gene transcription^5^. Whether this binding then leads to enhancer activation and nearby gene transcription depends on complex interactions with cofactors determined by the local motif composition^6^.

Dysregulation of GR levels and GC secretion are hallmarks of chronic stress-induced conditions, including depression, but also neurodegenerative diseases^7–12^. For example, in patients with post-traumatic stress disorder, hypersensitive GR and associated reductions in GC levels have been repeatedly found^13^. Besides implications in neuropsychiatric diseases, altered GC levels play a major role in the progression and metastasis of various types of cancer, Cushing syndrome, and metabolic disease^14^.

Treatment strategies have already been explored that attenuate GR signaling. For instance, in patients with psychotic depression, treatment with the clinically available but non-selective GR antagonist mifepristone (MIF) showed promising results, such as rapidly improving depression symptoms^15,16^. However, binding of MIF to the progesterone and androgen receptor and long-term high dosage requirements are considerable disadvantages^14^. In rodents, administration of MIF has proven beneficial to mitigate stress pathology and has been used to study the implication of GR in the dysregulation of the stress response^15,17^. Despite such effects, the outcome of MIF is difficult to interpret, as the MIF-GR complex recruits transcriptional coregulators and may show (partial) agonism^18,19^. An additional complication is its highly different plasma half-life between rodents and humans^20,21^.

To address the need to better target aberrant GR activation in various disorders, recent research has explored GR antagonists and mixed antagonists/agonists, such as the selective modulator CORT113176. Application of CORT113176 in a rat model of Alzheimer’s disease showed improvement of some cognitive deficits, which was not observed with MIF^22^. Partial agonism or selective antagonism arguably have advantages in clinical settings, as they potentially allow for disease-tailored intervention in the function of the ubiquitously expressed and vital GR. Yet, they can - at the same time - also complicate the interpretation of obtained results. An alternative yet unexplored solution to overcome above mentioned obstacles would be sustained depletion of GR at the protein level. Such depletion could circumvent partial agonism and inverse agonism of some selective inhibitors, avoid crosstalk with other types of receptors and prevent adaptations often encountered in genetic deletion approaches.

In the current study, we made use of proteolysis targeting chimeras (PROTAC) technology to deplete GR protein via the cell’s internal proteasome machinery. PROTACs, a new class of small molecules with promising translational potential for drug development, enable direct manipulation at the protein level by selectively inducing protein degradation^23,24^. They can be designed for any target protein to effectively induce degradation at nanomolar dosage by hijacking the ubiquitin-proteasome system (UPS). PROTACs perpetuate the degradation signal from one target protein to the next without being consumed in the process themselves. Hence, they provide an attractive approach to modulating the levels of target proteins via an event-driven mode of action rather than inhibition. Due to the low dose requirements, these drugs have an outstanding safety profile, making them interesting candidates for translational approaches^25^.

Here, we describe the successful development of a cereblon (CRBN)-recruiting PROTAC directed against the GR, and we compare its performance to existing pharmacological inhibitors in various model systems. Our results provide an in-depth functional characterization of a novel synthetic small-molecule for GR-degradation and demonstrate excellent performance in passively preventing ligand-induced gene expression activation in the absence of partial agonistic transcriptional triggering and crosstalk inhibition, allowing a straightforward interpretation of treatment outcomes. Lastly, we showcase a proof-of-concept application for targeted in-vivo GR depletion without the need for laborious and time-consuming intercrosses of genetically modified mouse lines and, hence, with direct relevance to disease models and translation potential for clinical use.

## Results

### GR-PROTAC design and linker optimization

To identify a CRBN-recruiting GR-PROTAC, we employed dexamethasone (DEX) as a small-molecule ligand for GR and lenalidomide for CRBN recruitment. By analyzing the crystal structure of DEX complexed with GR (protein data bank (PDB): 4UDC), we assessed the exit vector on the deeply embedded DEX and recognized a bile acid co-crystallized on the surface of GR at the entrance of the substrate-binding pocket (Fig. 1a)^26^. We hypothesized the bile acid could serve as a surrogate for a CRBN-recruiting ligand.

**Figure. 1:**
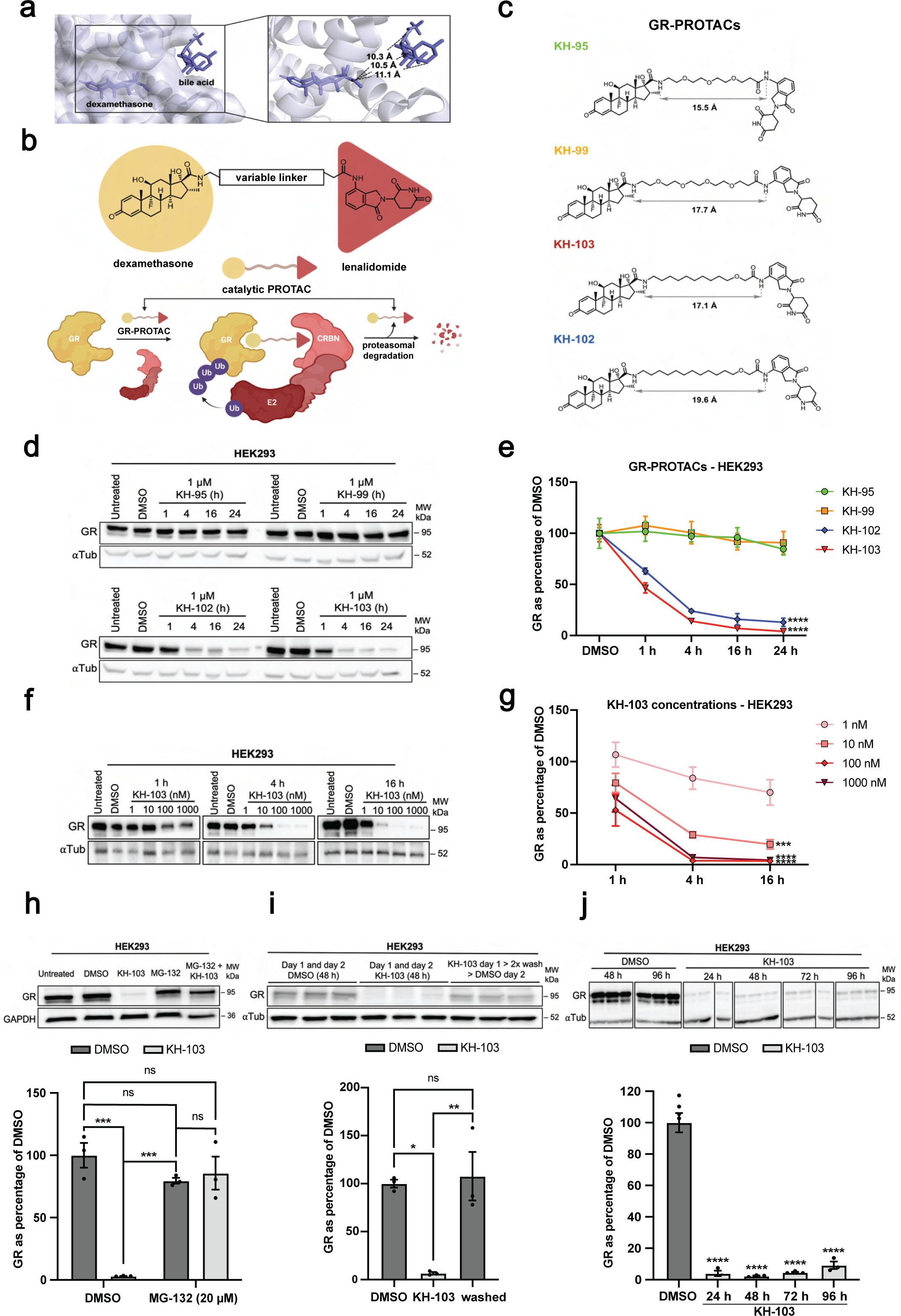
KH-103 and KH-102 were identified as two novel potent GR degraders. **a** The crystal structure of DEX and GR in a complex with bile acid (PDB: 4UDC) was used for the surrogate approach. **b** Schematic representation for the design of the GR-PROTACs (upper row) and the catalytic mode of action of PROTACs (lower row). **c** Four GR-PROTAC candidates and their individual linker lengths. **d** Representative immunoblot of GR in HEK293 cells following GR-PROTAC treatments at multiple time points. **e** Quantification of the immunoblots shown in d (N = 4). Two-way ANOVA showed significant main effects for time (F(4, 60) = 34.73, p = < 0.0001) as well as for GR PRTOACs (F(3, 60) = 106.10, p = < 0.0001). It also showed a significant interaction between both factors (F(12, 60) = 7.81, p = < 0.0001). Follow-up Dunnett’s multiple comparisons between compounds vs. DMSO at 24 h showed significant differences for both KH-102 (p = < 0.0001) and KH-103 (p = < 0.0001)). **f** Representative immunoblot of HEK cells treated with KH-103 at various concentrations. **g** Quantification of immunoblots shown in f (N = 3). Two-way ANOVA showed significant main effects for concentration (F(4, 30) = 22.91, p = < 0.0001) as well as for time (F(2, 30) = 14.43, p = < 0.0001). There was no significant interaction between both factors (F(8, 30) = 1.18, P = 0.3420). Follow-up Dunnett’s multiple comparison tests between various concentrations vs. 16 h time point showed significant differences for 10 nM (p = 0.0006), 100 nM (p = < 0.0001), and 1000 nM (p = < 0.0001). **h** Representative immunoblot and quantifications of GR degradation by KH-103 in presence of MG-132 (N = 3). Two-way ANOVA showed significant main effects for the MG-132 (F(1, 8) = 13.90, p = 0.0058) and for the KH-103 (F(1, 8) = 29.70, p = 0.0006). There was also a significant interaction between both (F(1, 8) = 38.20, p = 0.0003). Follow-up Tukey’s multiple comparison tests showed significant differences for DMSO/DMSO vs. DMSO/KH-103 (p = 0.0002), and DMSO/KH-103 vs. MG-132/DMSO (p = 0.0009). **i** Representative immunoblot and band quantification of GR in HEK293 cells washed from KH-103 by medium exchange (N = 3). Ordinary one-way ANOVA showed a significant difference (F(2, 6) = 14.40, p = 0.0051). Follow-up Tukey’s multiple comparisons showed significant differences for DMSO vs. KH-103 (p = 0.0102) and KH-103 vs. washed (p = 0.0070). **j** Representative immunoblot and quantification of GR in HEK293 cells treated with KH-103 for up to four days. N = 6 for DMSO (pooled 48 h and 96 h DMSO), and N = 3 for all other conditions. Ordinary one-way ANOVA showed a significant difference (F(4, 13) = 100.00, p = < 0.0001). Follow-up Dunnett’s multiple comparisons showed significant differences for DMSO vs. 24 h (p = < 0.0001), 48 h (p = < 0.0001), 72 h (p = < 0.0001), and 96 h (p = < 0.0001). P-values * < 0.05, ** < 0.01, *** < 0.001, **** < 0.0001.

Thus, the minimal linker length required for a functional GR-PROTAC could be anticipated by measuring the distance between the DEX exit vector and the bile acid on the surface. According to the measured distances, we set out and synthesized DEX-conjugated analogs with lenalidomide and varying linker lengths and composition (Fig. 1b). Four candidates were synthesized, two of which consisted of poly(ethylene glycol) (PEG) linkers (KH-95 and KH-99), while the other two were composed of alkyl linkers (KH-103 and KH-102) (Fig. 1c and supplementary Fig. 1 and 2).

Initial treatments of the human embryonic kidney 293 (HEK293) cells with 1 μM of the candidates revealed that KH-102 and KH-103 led to efficient and rapid degradation of GR cells within less than an hour, while KH-95 and KH-99 did not induce any degradation as determined by western blot (Fig. 1d, e). Roughly half of the GR proteins were depleted within the first hour of treatment, reaching an almost complete GR depletion from 16 hours (h) onwards. Since both GR-PROTACs connected by PEG linkers (KH-95 and KH-99) were inactive, linker composition showed to be crucial for PROTAC-mediated GR degradation. Interestingly, both PROTACs based on alkyl linkers designed by our surrogate approach resulted in efficient GR-PROTACs (KH-103 and KH-102).

We selected KH-103 as the most potent candidate for the treatment of cells with various concentrations of this compound and found that highly efficient degradation could still be achieved at a hundred times lower concentrations (10 nM), reaching almost full depletion at 100 nM (Fig. 1f, g). We then tested the proteasomal dependency of the KH-103 mechanism of degradation in HEK293 cells. As expected, GR depletion by KH-103 was inhibited in the presence of 20 μM of the proteasomal inhibitor MG-132 (Fig. 1h). Additionally, the removal of KH-103 by two medium exchanges showed an almost full recovery of GR within 24 h (Fig. 1i), indicating reversibility. When the medium was not exchanged, GR depletion was sustained up to 96 h (Fig. 1j).

### KH-103 shows efficient reversible degradation in various in-vitro models across tissue origins and species

We anticipated that KH-103 would degrade both human and rodent GR because the ligand binding site is conserved. Yet, it has been reported that PROTAC efficiency is cell-type and tissue-dependent, limited both by protein resynthesis rate and CRBN abundance^27^. Therefore, we assessed GR depletion in different cell types and across species. In the neuroblastoma mouse cell line, neuro 2a (N2a), similarly to HEK293, 100 nM of KH-103 was sufficient to induce almost complete degradation of GR (Fig. 2a-d). Likewise, we observed the same effect in a cell line expressing high levels of endogenous GR, the human lung carcinoma cell line A549 (Fig. 2e). N2a cells showed full maintenance of GR depletion by KH-103 up to 96 h (Fig. 2f). Also, upon medium exchange, N2a cells showed full recovery of GR protein levels (Fig. 1g). Efficient GR degradation was also achieved in mouse hippocampal and cortical primary cultures. This was confirmed by both immunoblotting (Fig. 2h) and immunofluorescence staining using a neuronal-specific marker, MAP2 (Fig. 2i), confirming KH-103’s broad applicability.

**Figure. 2:**
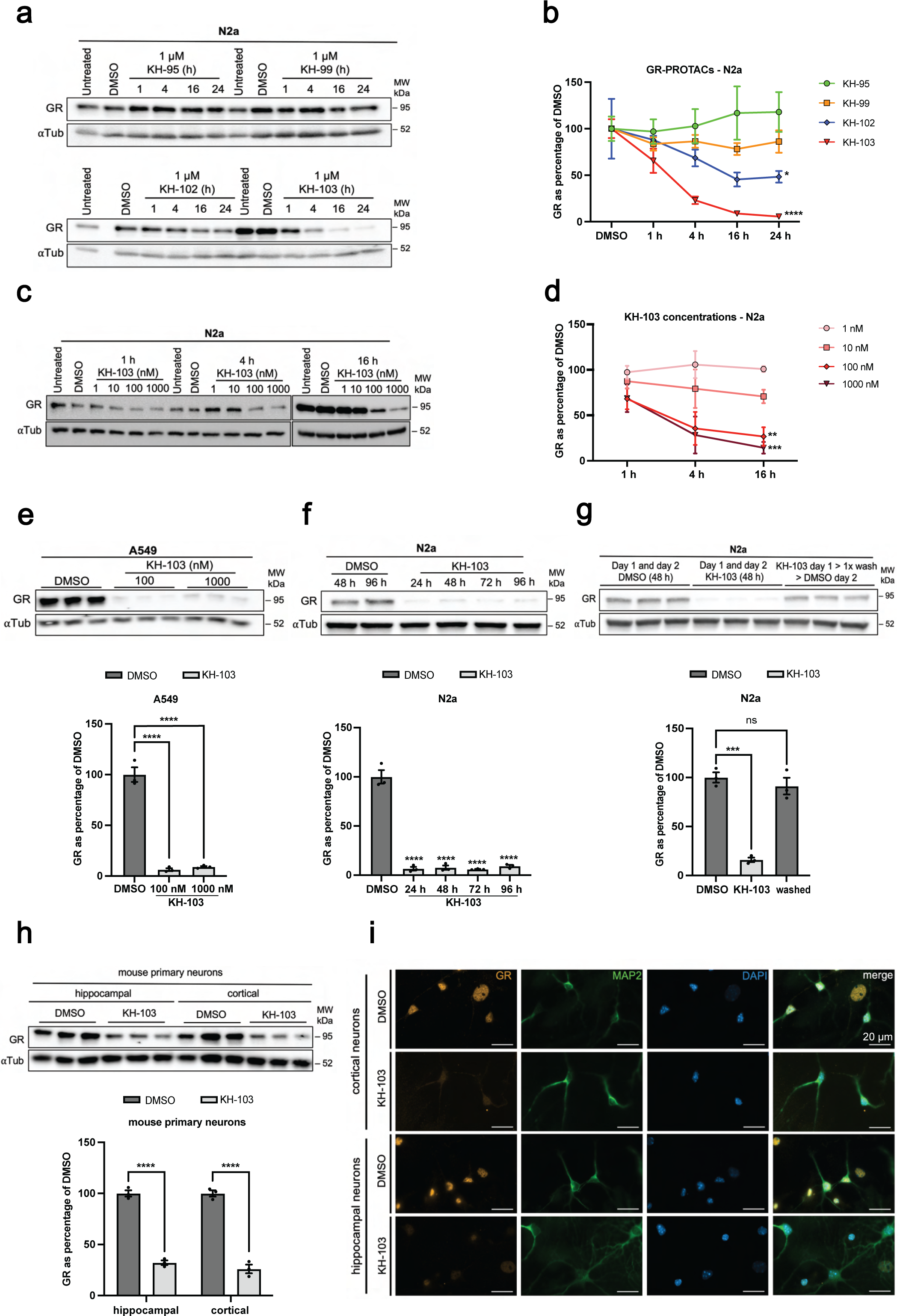
KH-103 induces degradation of GR in various cell types across species. **a** Representative immunoblot of GR in N2a cells following GR-PROTAC treatments at multiple time points. **b** Band quantification of the immunoblots shown in a. Two-way ANOVA showed significant main effects for time (F(4, 40) = 5.21, p = 0.0018) as well as for GR PRTOACs (F(3, 40) = 20.84, p = < 0.0001). It also showed a significant interaction between both factors (F(12, 40) = 2.51, p = 0.0145). Follow-up Dunnett’s multiple comparisons between compounds vs. DMSO at 24 h showed significant differences for both KH-102 (p = 0.0382) and KH-103 (p = < 0.0001)). **c** Representative immunoblot of comparison of treatment with KH-103 at various concentrations. **d** Quantification of immunoblots shown in c. Two-way ANOVA showed significant main effects for concentration (F(4, 30) = 14.21, p = < 0.0001) and a trend for time (F(2, 30) = 3.13, p = 0.0583). There was no significant interaction between both factors (F(8, 30) = 0.94, p = 0.4974). Follow-up Dunnett’s multiple comparison tests between various concentrations vs. 16 h time point showed significant differences for 100 nM (p = 0.0032) and 1000 nM (p = 0.0006). **e** Representative immunoblot and band quantification of KH-103 treatment in A549 cells. Ordinary one-way ANOVA showed a significant difference (F(2, 6) = 158.8, p = < 0.0001). Follow-up Tukey’s multiple comparisons showed significant differences for DMSO vs. 100 nM (p = < 0.0001) and 1000 nM (p = < 0.0001). **f** Representative immunoblot and band quantification of GR in N2a cells treated with KH-103 for up to four days (DMSO treatment for 96 h, GRs in all conditions are shown as a percentage of 48 h DMSO). Ordinary one-way ANOVA showed a significant difference (F(4, 10) = 150.70, p = < 0.0001). Follow-up Dunnett’s multiple comparisons showed significant differences for DMSO vs. 24 h (p = < 0.0001), 48 h (p = < 0.0001), 72 h (p = < 0.0001), and 96 h (p = < 0.0001). **g** Representative immunoblot and band quantification of GR in N2a cells washed from KH-103 by the medium exchange. Ordinary one-way ANOVA showed a significant difference (F(2, 6) = 60.32, p = 0.0001). Follow-up Dunnett’s multiple comparisons showed a significant difference for DMSO vs. KH-103 (p = 0.0001) and no significant difference between DMSO vs. washed (p = 0.5117). **h** Immunoblot and band quantification of GR in hippocampal and cortical primary mouse neurons. Two-way ANOVA showed significant main effects for KH-103 (F(1, 8) = 479.20, p = < 0.0001). There was no significant main effect for cell type (hippocampal or cortical) (F(1, 8) = 0.91, p = 0.3690) and no significant interaction between both factors (F(1, 8) = 0.91, p = 0.3690). Follow-up Sidak’s multiple comparisons between DMSO vs. KH-103 showed significant differences in both hippocampal (p = < 0.0001) and cortical (p = < 0.0001) primary mouse neurons. **i** Immunofluorescent staining of GR and MAP2, a neuronal marker, in hippocampal and cortical primary mouse neurons treated with KH-103. Scale bars are 20 μm. N = 3. P-values * < 0.05, ** < 0.01, *** < 0.001, **** < 0.0001.

### KH-103 induces nuclear translocation without triggering GR-mediated gene activation

DEX binding to GR is sufficient to cause conformational changes, which expose the nuclear translocation signal motif on the GR, causing GR translocation to the nucleus^28^. As KH-103 was designed based on the DEX binding site, we assessed whether binding of KH-103 to GR would also trigger GR translocation to the nucleus. Since our initial experiments (Fig. 1e) revealed that about half of the GR is already degraded within the first hour, we visualized the GR protein signal at various time points, starting already at 30 minutes (min) after treatment with KH-103 or DEX. Indeed, in line with immunoblot analysis, staining revealed a progressive loss of GR signal over time upon the addition of KH-103 both in HEK293 cells (Fig. 3a) and in A549 cells (Fig. 3b). DEX and KH-103 treated cells showed almost an exclusive nuclear signal for GR, indicating translocation of GR to the nucleus within 30 min. While the nuclear translocation was maintained during the 20 h of DEX treatment, the nuclear signal strongly faded over time after the addition of KH-103 (Fig. 3a, b). This observation indicated that either 1) the binding of KH-103 to GR has exposed the nuclear localization signal and induced its translocation from cytosol to the nucleus similarly to DEX, or 2) that the GR depletion occurred at different speeds in the cytosolic and nuclear UPS. To rule out that the full nuclear localization of GR was due to fast cytosolic GR degradation within 30 min, we repeated the staining in HEK293 cells at the 30 min time point in the presence of proteasome inhibitor MG-132. Again, DEX, KH-103, and KH-103 treated cells pre-incubated with MG-132 showed primarily nuclear signal for GR compared to primarily cytosolic GR signal in dimethyl sulfoxide (DMSO) and MG-132 control conditions (Fig. 3c), confirming rapid GR translocation to the nucleus by KH-103. This finding corroborates that the rapid and high GR degradation efficiency is primarily exerted by the nuclear UPS. This feature seems especially relevant for the depletion of nuclear proteins and transcription factors by PROTACs in general.

**Figure. 3:**
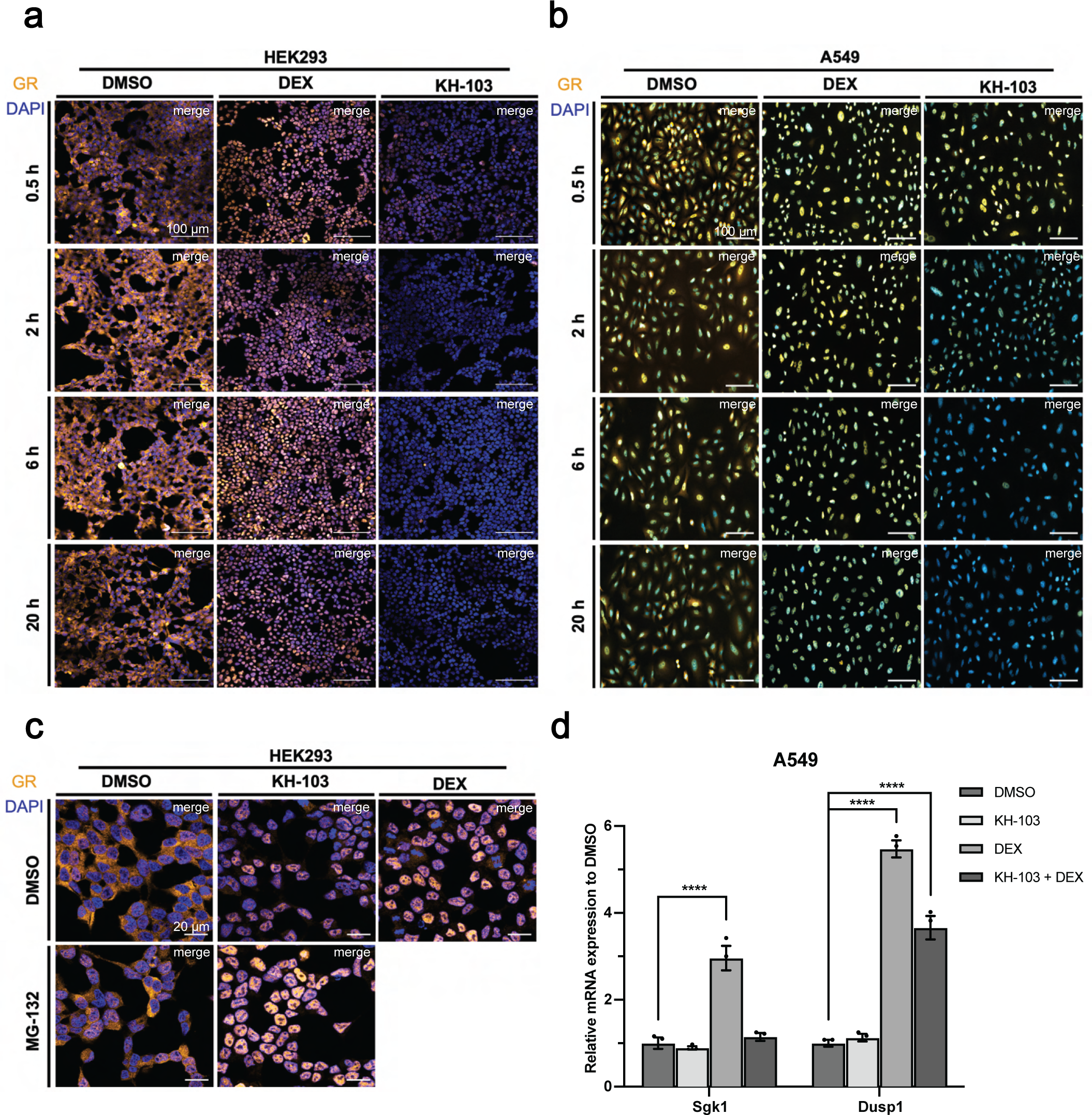
KH-103-mediated nuclear translocation of GR. Immunofluorescent staining of GR in HEK293 (**a**) and A549 (**b**) cells treated with DEX or KH-103 for several durations. Scale bars are 100 μm. **c** Co-treatment of KH-103 with MG-132 in HEK293 cells at 30 min. Scale bars are 20 μm. **d** Relative mRNA expression of Sgk1 and Dusp1 in A549 cells treated with DEX, or KH-103 or DEX and KH-103 for 2 h (N = 3). For Sgk1, two-way ANOVA showed a significant main effect for KH-103 (F (1, 8) = 34.77, p = 0.0004) and DEX (F (1, 8) = 45.92, p = 0.0001). There was also significant interaction between KH-103 and DEX (F(1, 8) = 27.47, p = 0.0008). Follow-up Dunnett’s multiple comparisons showed a significant difference between DMSO vs. DEX (p = < 0.0001). For Dusp1, two-way ANOVA showed significant main effects for KH-103 (F (1, 8) = 22.72, p = 0.0014) and DEX (F (1, 8) = 391.40, p = < 0.0001). There was also a significant interaction between KH-103 and DEX (F(1, 8) = 30.06, p = 0.0006). Follow-up Dunnett’s multiple comparisons showed significant differences between DMSO vs. DEX (p = < 0.0001) and DMSO vs. KH-103 + DEX (p = < 0.0001). P-values **** < 0.0001.

The initial translocation of GR to the nucleus upon KH-103 raised the possibility that KH-103 treatment might trigger GR-dependent transcriptional activity. Therefore, we explored this possibility experimentally.

Following 2 h of treating A549 cells with KH-103 or DEX, we assessed the relative mRNA expression of two well-known GR targets, the immediate early genes, the serum/glucocorticoid regulated kinase 1 (Sgk1) and the dual-specificity phosphatase 1 (Dusp1). The reverse transcription-quantitative polymerase chain reaction (RT-qPCR) results showed that DEX treatment, but not KH-103 treatment, significantly increased expression (Fig. 3d)^29,30^.

Moreover, co-treatment of KH-103 and DEX shows a dampened Sgk1 and Dusp1 increase as compared to DEX alone (Fig. 3d). These results point towards a potential competition between depletion and transcriptional regulation due to GR’s shared binding site for its natural ligands and KH-103. On the one hand, this resulted in reduced GR activation, as observed above (Fig. 3d); on the other hand, it might also prevent efficient GR degradation. A similar competition is expected in-vivo; therefore, it seemed particularly important to further assess KH-103 degradation characteristics in the co-presence of GR ligand.

### KH-103 degrades GR despite competition with DEX/cortisol binding

To test whether a putative competition would lead to decreased GR degradation efficiency, we co-treated HEK293 cells with KH-103 and with either 100 nM DEX or with a high-end physiological level (100 nM; 20x dissociation constant (K_D_)) of the GR-ligand cortisol. We predicted less competition with cortisol due to the lower binding affinity and shorter dissociation time of cortisol from GR in comparison to DEX^31^. Immunofluorescent staining after 12 h showed almost complete degradation in co-presence with cortisol but some remaining GR protein in KH-103 co-treatment with DEX (Fig. 4a). This confirms a competition between the synthetic activating ligand DEX and KH-103. Such competition appeared negligible for the endogenous ligand cortisol/corticosterone at the tested molar ratios. Thus, GR depletion should be feasible in the presence of physiological concentrations of cortisol/corticosterone. We further assessed whether 1 h of pretreatment with DEX would compromise efficient depletion upon KH-103 addition. Prior activation of GR by DEX slightly reduces the efficiency, but importantly it does not prevent the following depletion of GR by KH-103 (Fig. 4b).

**Figure. 4:**
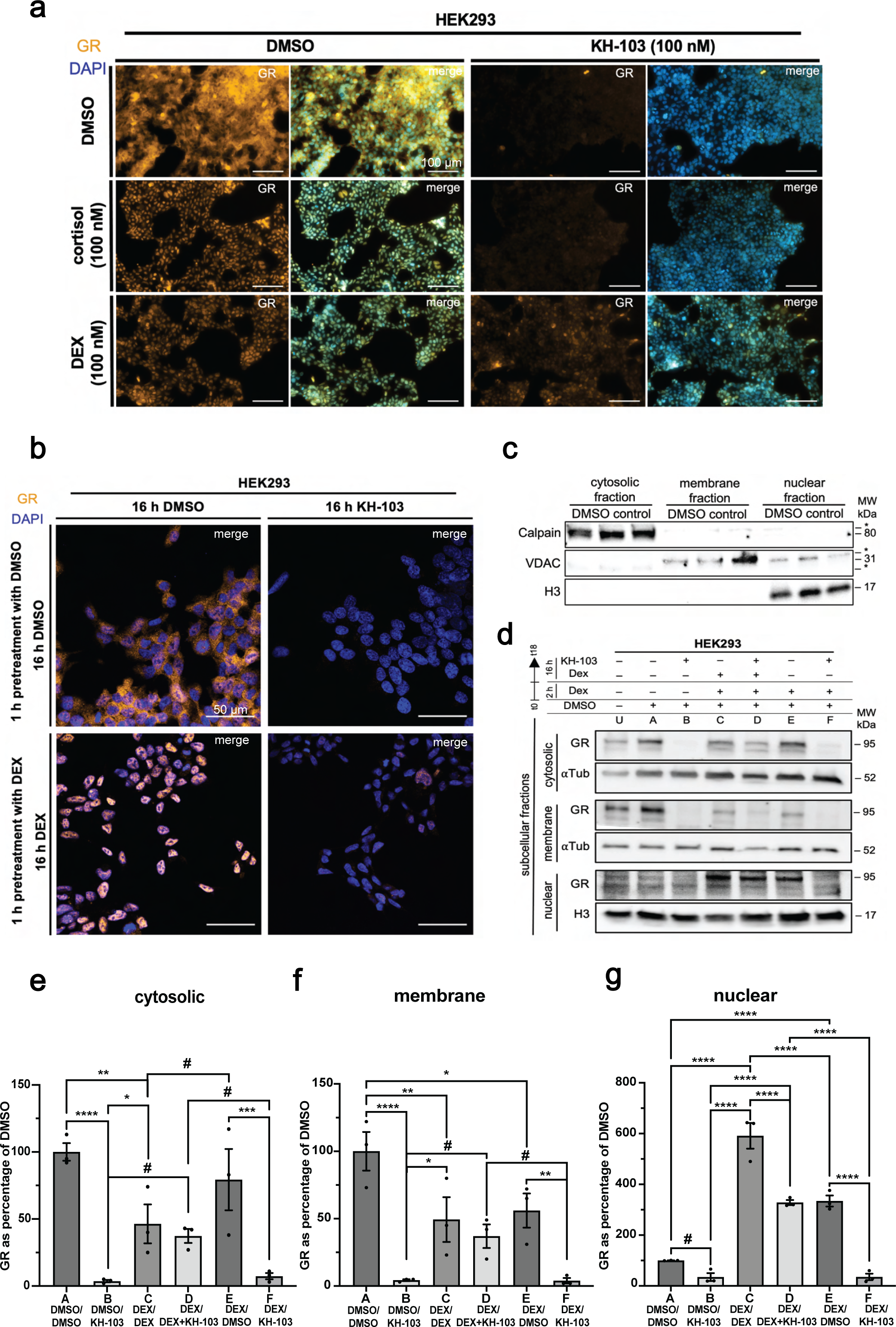
Assesment of KH-103 efficiency in co-presence with DEX. **a** Representative immunofluorescent staining of GR in HEK293 cells treated with KH-103, or DEX, or cortisol, or DEX + KH-103, or cortisol + KH-103. Scale bars are 100 μm. **b** Representative immunofluorescent staining of GR in HEK293 cells that were pretreated with DEX for 1 h followed by KH-103 treatment. Scale bars are 50 µm. **c** Representative immunoblot of fraction-specific markers in cytosolic, membrane, and nuclear fractions obtained from HEK293 cells. *unspecific bands. **d** Representative immunoblots of GR in cytosolic, membrane, and nuclear fractions obtained from HEK293 cells treated under various conditions. U: untreated, A: DMSO, B: 2 h pretreatment with DMSO followed by 16 h treatment with KH-103 (DMSO/KH-103), C: DEX/DEX, D: DEX/DEX+KH-103, E: DEX/DMSO, F: DEX/KH-103. E and F conditions included an extra wash step after the pretreatment with DEX. **e** Band quantification of GR in the cytosolic fraction. Two-way ANOVA showed a significant main effect for KH-103 (F(1, 12) = 38.84, p = < 0.0001) and a significant interaction between KH-103 and DEX (F(2, 12) = 7.53, p = 0.0076). There was no significant main effect for DEX (F(2, 12) = 0.43, p = 0.6595). **f** Band quantification of GR in the membrane fraction. Two-way ANOVA showed a significant main effect for KH-103 (F(1, 12) = 35.41, p = < 0.0001) and a significant interaction between KH-103 and DEX (F(2, 12) = 7.21, p = 0.0088). There was no significant main effect for DEX (F(2, 12) = 2.06, p = 0.1698). **g** Band quantification of GR in the nuclear fraction. Two-way ANOVA showed a significant main effect for KH-103 (F(1, 11) = 106.30, p = < 0.0001) and DEX (F(2, 11) = 129.20, p = < 0.0001). There was also a significant interaction between KH-103 and DEX (F(2, 11) = 9.88, p = 0.0035). Follow-up Fisher’s LSD tests for all three fractions are summarized in supplementary tables 1-3. N = 3. P-values # < 0.1, * < 0.05, ** < 0.01, *** < 0.001, **** < 0.0001.

In addition to the mere binding competition, ligand-dependent activation also induces changes in GR cellular relocalization, which could also potentially prevent GR depletion by KH-103. To test whether depletion efficiency varies according to cellular localization, we performed a fractionation experiment and assessed GR levels upon DEX versus no DEX in cytoplasmic, nuclear, and organelle membrane-bound fractions. Comparing compartment-specific markers, Calpain (cytosol), voltage-dependent anion channel (VDAC) (organelle-membrane), and Histone 3 (H3) (nuclear) showed enrichment of these proteins, thus high purity of the fractions (Fig. 4c). Immunoblotting analysis of each fraction revealed significant interactions between DEX and KH-103 in each cellular fraction (Fig. 4d-g). Indeed, KH-103 is highly efficient at inducing a significant reduction in GR of all three cytoplasmic, organelle-bound membrane, and nuclear fractions in comparison to DMSO controls (Fig. 4e-g, conditions A vs. B). In line with immunostainings, upon co-treatment with DEX, the efficiency of KH-103 in degrading GR is reduced in all three fractions (C vs. D). Upon washing the cells after 2 h of pretreatment with DEX and incubating them with KH-103, the efficiency is recovered (Fig. 4e-g, B: vs. D: vs. F:). DEX treatment significantly reduced GR protein, both in cytosolic and membrane fractions, compared to the DMSO control, due to the expected ligand-induced translocation to the nucleus. Accordingly, GR protein significantly increased in the nuclear fraction following DEX due to the DEX-induced translocation (Fig. 4g A: vs. C:).

Interestingly, there is a portion of GR in both cytosolic and membrane fractions that, after treatment with DEX, did not translocate to the nucleus and is also not degraded by KH-103 (Fig. 4e, f C: vs. D:). The nuclear GR containing the DEX activated/translocated GR was significantly reduced when KH-103 was added (Fig. 4g C: vs. D:) implying once more that KH-103 can also degrade ligand-activated nuclear GR.

Medium wash after 2 h of DEX pretreatment decreased GR levels significantly in the nuclear fraction (Fig. 4g C: vs. E:), likely indicating relocation to the cytoplasm. Combined 16 h KH-103 and DEX treatment following 2 h DEX significantly decreased GR levels in comparison to continuous 18 h DEX treatment (Fig. 4g C: vs. D:), whereas washing out DEX after 2 h enhances depletion significantly (Fig. 4g D: vs. F:). Nevertheless, medium wash after DEX pretreatment did not fully return GR levels back to baseline (Fig. 4g A: vs. E:), indicating incomplete elimination of DEX molecules from the cells, e.g., those bound to GR. This is in agreement with the very slow DEX washout kinetics observed previously^32^. Furthermore, we newly observed that a portion of GR in the presence of DEX (Fig. 4e, C: vs. D:) remains in the cytoplasm and resists degradation by KH-103, raising the question of whether KH-103 might preferentially degrade some isoforms of GR while maybe having limited access to other GR isoforms. Therefore, we next investigated which GR isoforms are degradable by KH-103.

### The presence of ligand-binding domain (LBD) is required for GR isoforms to be degraded by PROTAC KH-103

The GR gene gives rise to different transcriptional isoforms with different and sometimes opposing functions (GR-**α**, GR-β, GR-□, GR-A, and GR-P). They differ in their C terminal domain, which is important for ligand binding. GR-β, GR-A, and GR-P lack a full LBD. Alternative translational initiation further generates translational isoforms (GR**α**-A, GR**α**-B, GR**α**-C1-C3, GR**α**-D1-D3) with varying truncation of the N terminal domain, which is important for interaction with co-regulatory factors. Here we selected two translational isoforms, GR**α**-C3 and GR**α**-D3. Because the KH-103 design relies on binding to the LBD, we expected that only isoforms with an intact LBD would be degradable. Therefore, we tested which isoforms of GR are degradable by KH-103 by cloning the cDNA of all the splice variant isoforms as well as the two selected translational isoforms (GR**α**-C3 and −D3) into degradation TAG (dTAG) plasmids^33^. The expressed cDNAs harbored two HA tags and a variant of FK506-binding Protein 12 (FKBP12^F36V^) that could be targeted by the dTAG13 compound. This served as an internal control for degradability. To test whether generally additional tagging would interfere with the tertiary complex formation, we transiently expressed GR tagged with an enhanced green fluorescent protein (EGFP). Despite the large size of the tag, we still observed efficient degradation upon KH-103 treatment (Fig. 5a). Separate and transient expression of different GR isoforms in dTAG plasmids showed the correct size for all constructs (Fig. 5b, c). Moreover, dTAG13 treatment in HEK293 cells degraded all isoforms, confirming that all the tested isoforms could be targeted by a CRBN-recruiting PROTAC for ubiquitination and depletion via the proteasomal pathway. Incubation with KH-103 revealed that both C and N terminally tagged GR-α as well as N-terminally tagged GR-□ and C terminally tagged GR-α-C3 and D3 showed degradation (Fig. 5d, e), consistent with the presence of a ligand binding domain in these isoforms. In contrast, KH-103 did not induce degradation of N-terminally tagged GR-β, GR-P, and GR-A isoforms, in line with the absence of an LBD in those isoforms. Interestingly, in the case of the GR-A isoform, we even observed a significant increase in GR-A levels after treatment with KH-103 in comparison to the DMSO control. Since GR-A was expressed exogenously, this could indicate that degradation of the endogenous isoforms, GR-α and GR-□, leads to the stabilization of GR-A proteins.

**Figure. 5:**
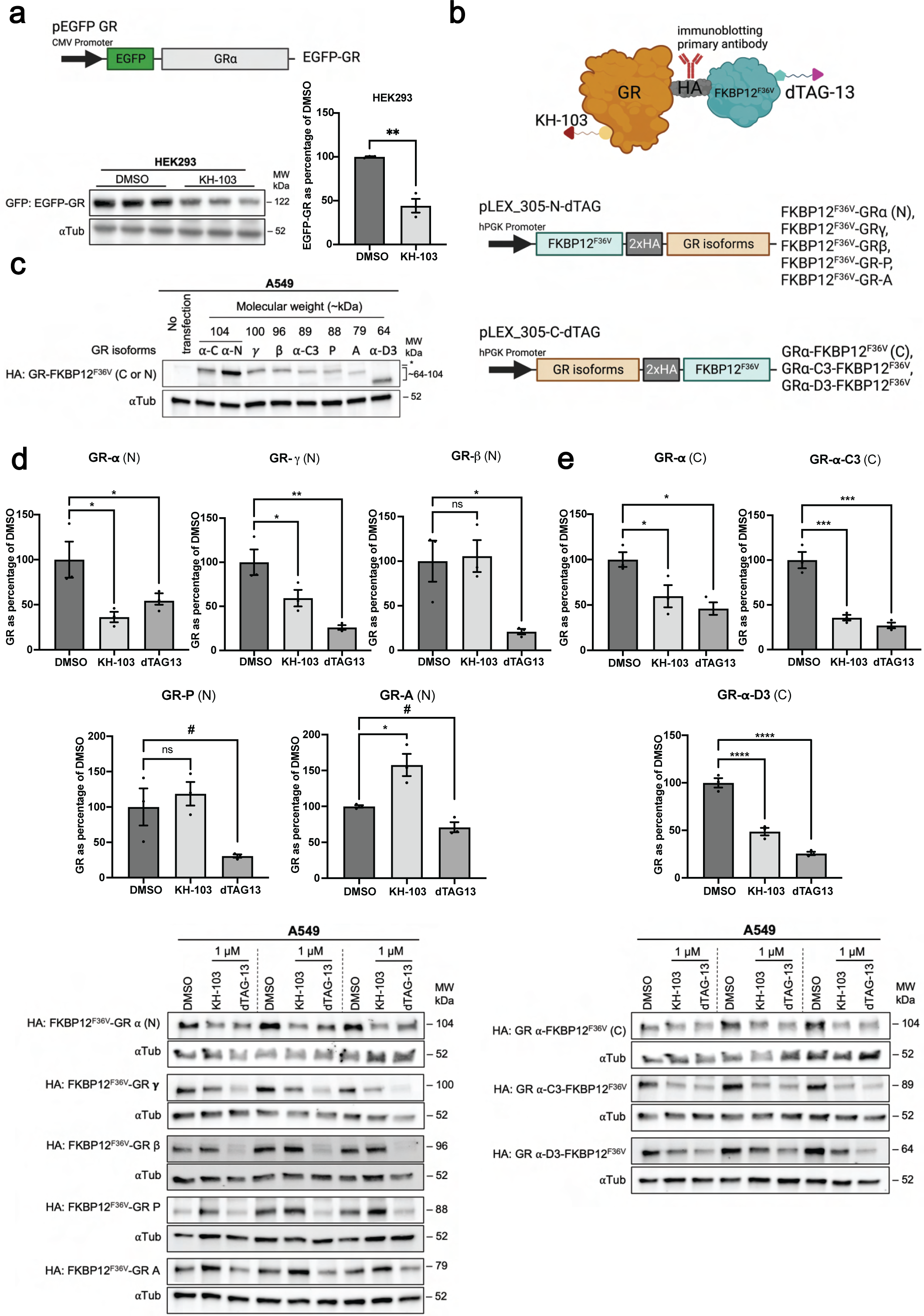
GR isoforms with intact LBD are degradable by KH-103. **a** Schematic depiction of plasmid used for expression of EGFP-GR fusion protein, immunoblot, and band quantification of HEK293 cells transiently expressing EGFP-GR and treated with KH-103. Unpaired two-tailed t-test showed a significant difference between DMSO and KH-103 (t(4) = 7.08, p = 0.0021). **b** Schematic depiction of GR isoforms in fusion with HA and FKBP12^F36V^ and their expression cassettes in dTAG plasmids. These proteins were targeted for degradation upon treatment with either KH-103 or dTAG13 and were detected by HA antibody. **c** Immunoblot of transiently expressed GR isoforms in A549 cells. *: unspecific band detected by HA antibody. **d** Band quantification and immunoblot of transcriptional GR isoforms transiently expressed in A549 cells treated with KH-103 or dTAG13. **e** Band quantification and immunoblot of translational GR isoforms transiently expressed in A549 cells treated with KH-103 or dTAG13. Results of ordinary one-way ANOVA and Holm-Sidak’s multiple comparison tests for transcriptional and translational isoforms are summarized in supplementary table 4. N = 3. P-values # < 0.1, * < 0.05, ** < 0.01, *** < 0.001, **** < 0.0001.

### KH-103’s unique characteristics complement current tools to study GR’s genomic actions

The finding that KH-103 degrades all transcriptionally active isoforms of GR encouraged us to assess how well it can counteract DEX-induced transcriptional changes. Despite preventing agonist binding, classical small-molecule antagonists allow the binding of GR to the DNA and, to varying degrees, interactions with transcriptional co-regulators. GR-PROTACs eliminate the protein and hence prevent any DNA interaction of GR and any co-regulator. To assess the functional consequences of such an alternative mode of action, we compared KH-103 to MIF, a non-selective GR antagonist with cross-reactivity with progesterone and, to a lesser extent, androgen receptors, and to CORT113176, a promising selective non-steroid GR modulator with partial agonistic actions, currently in a phase 2 trial for Amyotrophic Lateral Sclerosis (ALS)^34^.

We cultured A495 cells and exposed them to different treatment regimens, including DEX (100 nM), MIF (1 μM), CORT113176 (1 μM), KH-103 (100 nM) or a combination thereof with varying exposure time and order (see treatment scheme Fig. 6a). We then assessed the consequences on gene expression using RNA sequencing. We were interested in three questions. The first question aimed at determining KH-103’s potential to block DEX-induced gene expression changes (Fig. 6a I). The second question assessed KH-103’s potential to reverse DEX-induced gene expression changes (Fig. 6a II). The third question investigated the transcriptional effects of KH-103 in comparison to other inhibitors in the absence of the ligand DEX (Fig. 6a III).

**Figure. 6:**
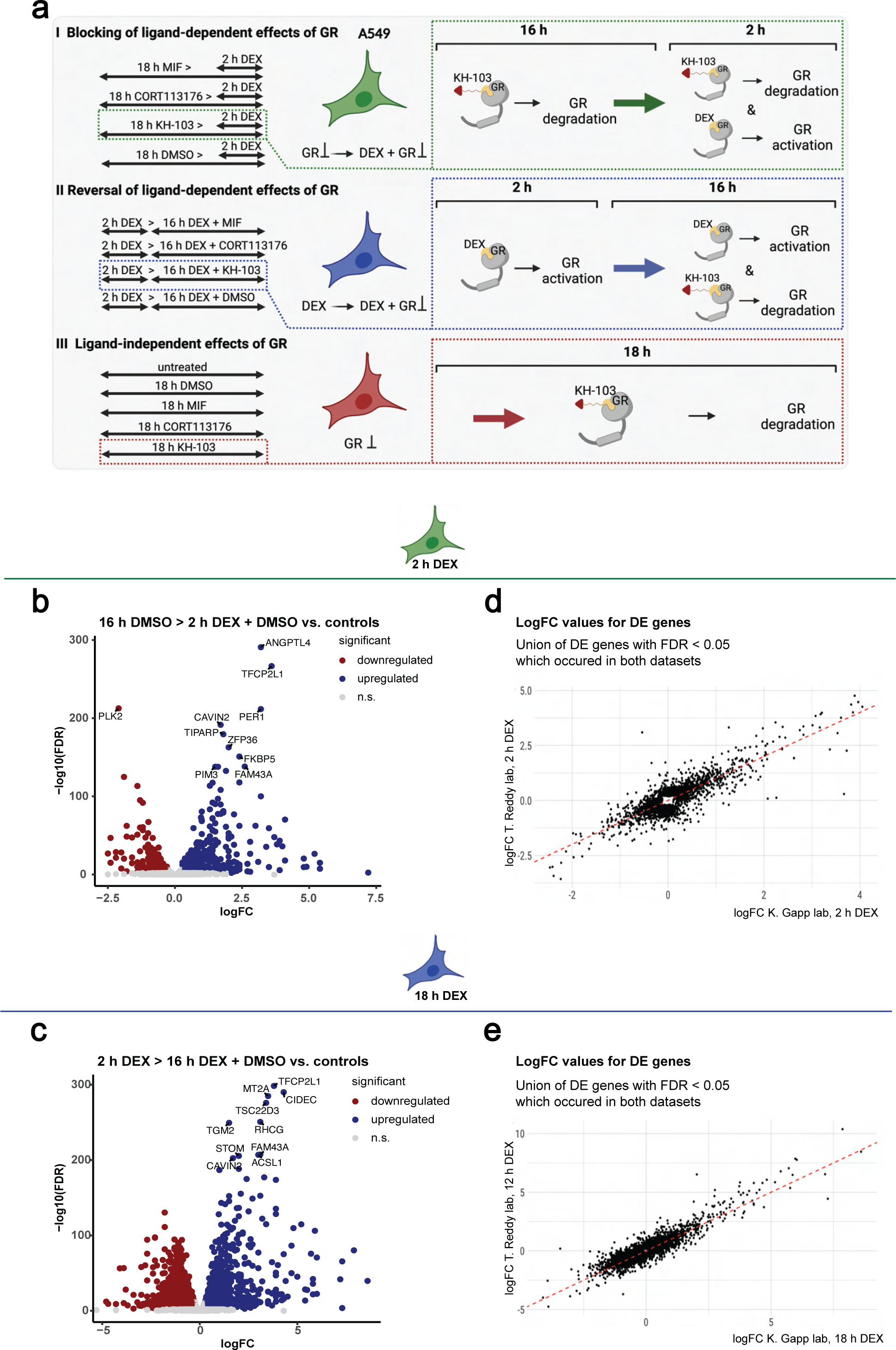
The DEGs we identified upon 2 h, and 18 h of DEX treatments in A549 cells align with the other existing RNAseq and ChIP-seq datasets. **a** Schematic depiction of the RNAseq experiment treatment groups in A549 cells. b Correlation of logFC values between DEGs with the false discovery rate (FDR) < 0.05 of T. Reddy lab after 2 h DEX treatment and K. Gapp lab 2 h DEX treatment. c Correlation of logFC values between DEGs with FDR < 0.05 of T. Reddy lab after 12 h of DEX treatment and K. Gapp lab 18 h DEX treatment. d Distribution of logFC values of DEGs after 2 h DEX treatment combined with GR binding information extracted from T. Reddy lab after 2 h DEX ChIP-sequencing. e Distribution of p-values of DEGs after 2 h DEX treatment combined with GR binding information extracted from T. Reddy lab after 2 h DEX treatment. f Distribution of logFC values of DEGs after 18 h DEX treatment combined with GR binding information extracted from T. Reddy lab after 12 h DEX treatment. g Distribution of p-values of DEGs of 18 h DEX dataset combined with GR binding information extracted from T. Reddy lab after 12 h DEX treatment. h LogFC of DEGs after 2 h DEX treatment plotted in relation to their TSS’ minimum distance to GR peaks, extracted from T. Reddy lab 2 h DEX treatment. i LogFC of DEGs after 18 h DEX treatment plotted in relation to their TSS’ minimum distance to GR peaks, extracted from T. Reddy lab 12 h DEX treatment.

We first confirmed that 2 h (Fig. 6a I, b) and 18 h (Fig. 6a II, c) DEX treatment elicited significant gene expression changes. To validate our experimental setup, we next compared our DEX-induced gene expression changes (2 h and 18 h DEX treatment versus DMSO) with a publicly available dataset of 2 h and 12 h DEX treatments in the same cell line, respectively. Direction and log fold changes (logFC) showed a high correlation between both datasets (Fig. 6d, e). Furthermore, integration with publicly available chromatin immunoprecipitation sequencing ((ChIP)-seq) data for GR showed a set of differentially expressed genes (DEGs) with clear GR peaks upon 2 h or 12 h DEX treatment, corroborating that these genes are likely directly regulated by GR (supplementary Fig. 4-5)^6^. Altogether, our benchmarking confirmed the reproducibility and quality of our transcriptomics data.

### KH-103 blocks DEX effects with increased efficiency compared to other inhibitors

To address question number one, the compounds’ potential to block DEX-induced gene expression changes, A549 cells were first exposed to 16 h of GR modulator/degrader (MIF, CORT113176, or KH-103) or DMSO as a control, followed by 2 h of simultaneous exposure to the modulator/degrader and DEX (Fig. 6a I). In comparison to MIF and CORT113176, depleting GR with KH-103 prior to DEX exposure resulted in enhanced blockage of DEX-induced transcriptional changes (Fig. 7a). This result was further corroborated by hierarchical clustering, depicting that KH-103 pretreatment segregates from a cluster formed by CORT113176 and MIF, apart from the expected segregation from DEX only and the control condition (supplementary Fig. 6).

**Figure. 7:**
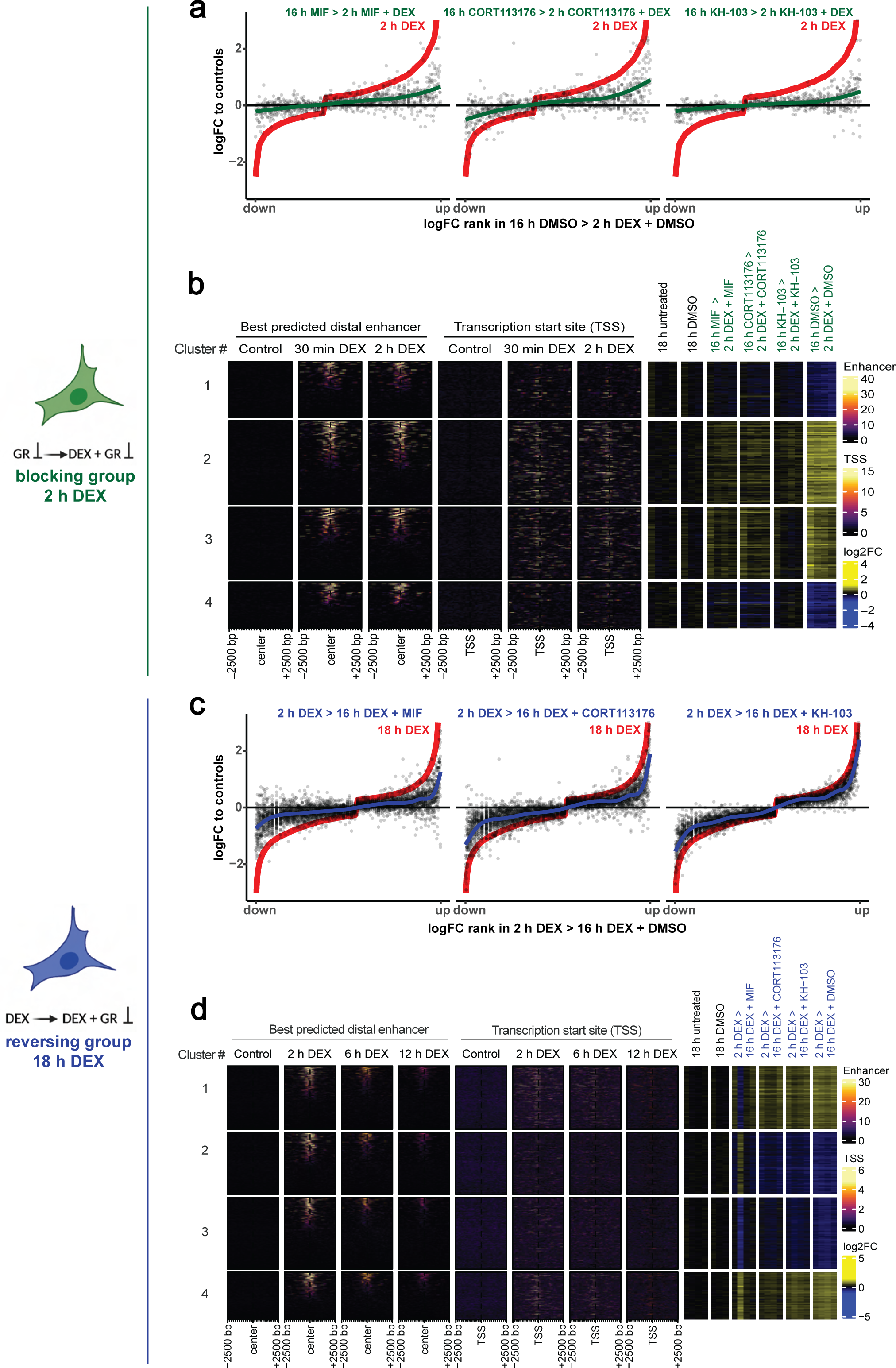
KH-103 efficiently blocks DEX-induced transcriptional changes. **a** Comparison of logFC of DEGs upon 16 h pretreatment with CORT113176, KH-103, or MIF prior to 2 h of DEX exposure and logFC of DEGs upon no pretreatment prior to the 2 h DEX condition (red line = 2 h DEX condition). A lack of inhibition would show as the black dots following the red line, while a full inhibition would show as the black dots around the x-axis. **b** Heatmap of DEGs of the blocking treatment groups (see Figure 6 a i) plotted aside GR binding information + / − 2500 base pairs (bp) centered around their TSS, as well as + / − 2500 bp centered around their best predicted distal enhancer, after 30 min and 2 h DEX treatment extracted from T. Reddy ChIP-sequencing data. **c** Comparison of logFC of DEGs upon 16 h treatment with inhibitors or KH-103 in co-presence of DEX after 2 h of DEX pretreatment, versus logFC of same genes upon 18 h DEX condition. **d** Heatmap of DEGs of the reversing treatment groups plotted aside GR binding information + / − 2500 bp centered around their TSS, as well as + / − 2500 bp centered around their best predicted distal enhancer, after 2 h, 6 h, and 12 h DEX treatment extracted from T. Reddy ChIP-seq data.

Furthermore, four k-means-based-clusters are segregated into two DEX-up-regulated and two DEX-down-regulated clusters. Additionally, this reveals slightly differential responsiveness to KH-103 across clusters, with cluster 2 being the least responsive despite more effective blockage compared to the inhibitors. Enhanced blockage by KH-103 is also evident in all other clusters but most evident in the up-regulated genes clusters: 2 and 3 (Fig. 7b). The genes up-regulated by DEX were less efficiently blocked by MIF and CORT113176 pre-incubation (clusters 2 and 3). These clusters showed more GR signal in the vicinity of their transcription start site (TSS), and about half showed GR binding at least at one putative distal enhancer^24^, suggesting direct regulation via GR binding. Clusters 1 and 4 are the genes that were down-regulated upon DEX treatment, with the least responsiveness to CORT113176 in cluster 4. Despite the presence of GR signal on distal enhancers for about half of these genes when selecting the putative enhancer with the highest GR signal, we observed fewer GR signals near their TSS compared to the up-regulated genes by DEX (clusters 2 and 3). These genes are thus potentially not regulated by GR binding to their promoter (Fig. 7b). Overall, irrespective of GR signal, the clustering of up-regulated genes showed which genes are sensitive to any GR antagonists, and the clustering of down-regulated genes revealed genes sensitive to CORT113176.

Comparison of the expression levels of individual DEX-induced genes that were efficiently blocked by KH-103 with their expression upon other inhibitors pretreatment revealed 100 genes that were not blocked by CORT113176 and MIF (supplementary Fig. 8a). In another subset of 6 DEX-induced genes, CORT113176 and MIF changed their expression significantly in the opposite direction of DEX, showing some inverse agonism, while KH-103 pretreatment efficiently prevented effects of DEX (WWTR1, MARCKS, SIX5, B4GALT5, FZD2, and IRAK2) resulting in no significant difference in expression compared to DMSO control (supplementary Fig. 8b). This again highlights the benefit of depletion of GR at the protein level as an effective passive approach to block DEX-induced transcriptional changes.

### KH-103 shows no inverse agonism

Addressing question number two, the potential of KH-103 to reverse DEX-induced gene expression changes, cells were exposed to 2 h of DEX alone followed by 16 h of simultaneous exposure to DEX and modulator/degrader or DEX and DMSO (Fig. 6a II). Likely due to the above-described competition between DEX and KH-103 (Fig. 4 & supplementary Fig. 3), reversal of DEX-induced changes by KH-103, after a 2 h DEX pretreatment, was overall less efficient than blocking of DEX-induced changes upon KH-103 pretreatment (Fig. 7c, d).

Nevertheless, KH-103 showed increased efficiency over the two inhibitors in the reversal of 44 genes (supplementary Fig. 9a). Moreover, for 19 genes, KH-103 treatment reversed their expression to a level indistinguishable from DMSO, while the inhibitors led to significant overcompensation of DEX-induced changes in the opposite direction, suggesting inverse agonism (supplementary Fig. 9b).

Inspection of the GR peaks at TSS revealed that DEX up-regulated genes (clusters 1 and 4) mostly showed GR ChIP-seq signal, yet not all of them showed GR at their enhancer and hence are not all conclusively direct GR targets (Fig. 7d). DEX-down-regulated genes also showed less GR signal at TSS yet some signal at a subfraction of enhancers, corroborating the findings from the above-mentioned blocking conditions that suggest indirect regulation or secondary effects of the recruitment of the transcriptional machinery to other sites. Based on the four k-means clusters, we saw both DEX-up and −down-regulated genes segregating into two clusters, one that shows more efficient reversal (cluster 3 and 4 respectively) and one reflecting less efficient reversal (cluster 1 and 2 respectively) by inhibitors, independent of GR signal at TSS or enhancers.

### KH-103 exerts “passive” inhibition’ and is specific

Our third question was whether, in the absence of ligand, GR inhibition per se is not expected to induce transcriptional changes, yet some intentional agonism of selective inhibitors respectively has been reported^35–37^. To address this third question, we profiled gene expression following 18 h treatment with either inhibitor or KH-103 in the absence of DEX (Fig. 8a, b). To this end, cells were exposed to 18 h of modulator/degrader (MIF or CORT113176 or KH-103) or vehicle (DMSO) (Fig. 6a III). Analyzing DEGs revealed thousands of significantly affected transcripts in MIF and CORT113176, consistent with partial agonistic properties. Such agonism can be explained by the unaffected DNA binding potential of inhibitor-bound GR that is absent in case of GR protein depletion (supplementary Fig. 6). KH-103 treatment merely affected the expression of 13 genes significantly (Fig. 8a, b). This indicates a passive mode of action. Using RT-qPCR, we confirmed that among those 13 DEGs in the KH-103 condition, only period circadian regulator 1 (Per1) and FK506-binding protein 51 (FKBP5) showed high fold-changes after 18 h of KH-103 treatment (supplementary Fig. 10).

**Figure. 8:**
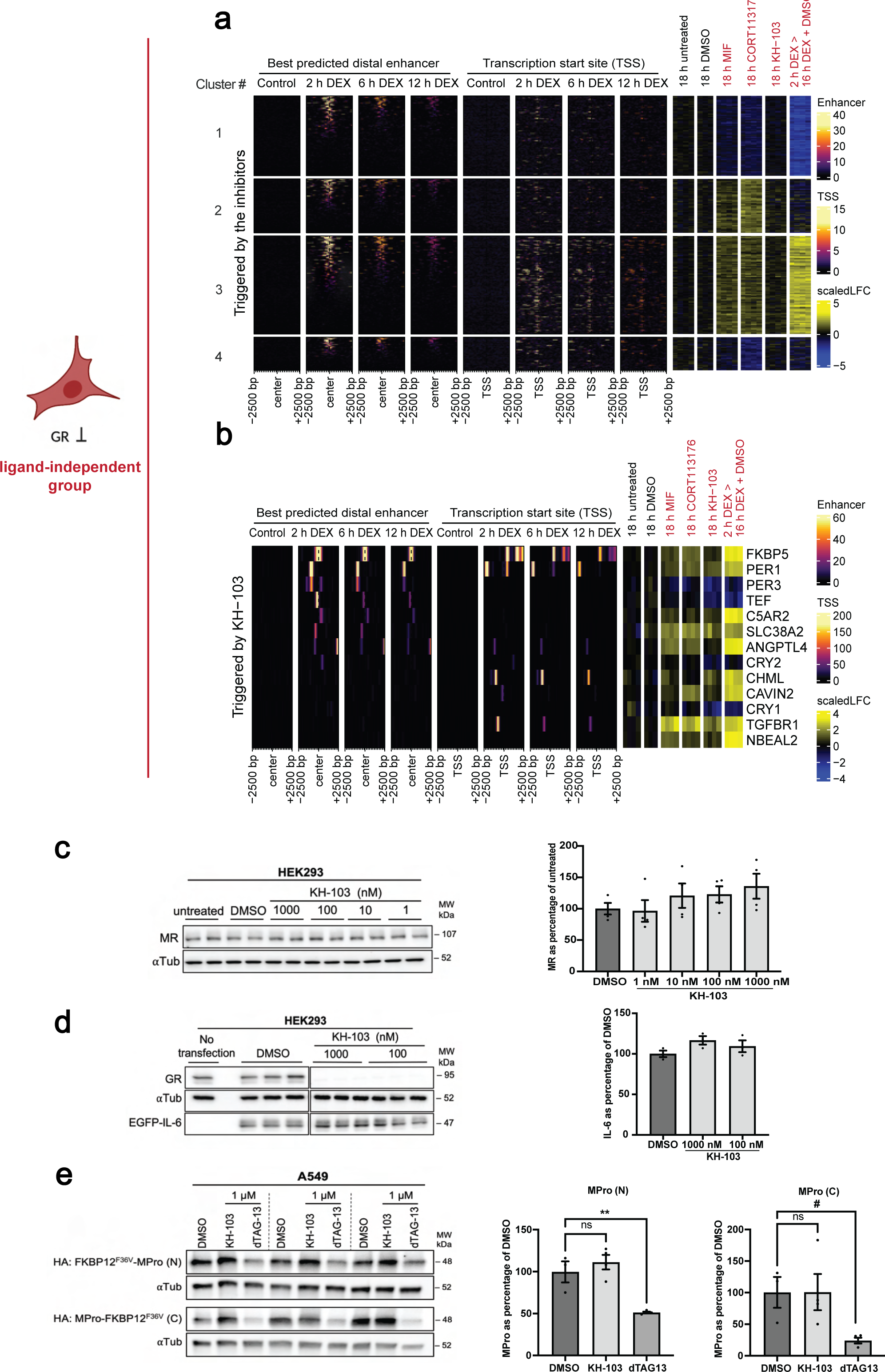
KH-103 treatment alone has no nonspecific transcriptional effects. **a** Heatmap of DEGs of the ligand-independent treatment groups plotted aside GR binding information + / − 2500 bp centered around their TSS, as well as + / − 2500 bp centered around their best predicted distal enhancer, after 2 h, 6 h, and 12 h DEX extracted from T. Reddy ChIP-seq data. **b** Heatmap of 13 DEGs triggered by KH-103 plotted aside GR binding information + / − 2500 bp centered around their TSS, as well as + / − 2500 bp centered around their best predicted distal enhancer, after 2 h, 6 h, and 12 h DEX treatment extracted from T. Reddy ChIP-seq datasets. **c** Representative immunoblot and band quantification of MR in HEK293 cells treated with KH-103 at multiple concentrations. N = 4 independent experiments. Ordinary one-way ANOVA showed no significant difference between DMSO and different KH-103 concentrations (F(4, 15) = 1.03, p = 0.4228). Follow-up Dunnett’s multiple comparisons tests also showed no significant differences. **d** Immunoblot and quantification of EGFP-IL6 transiently expressed in HEK293 cells treated with KH-103. N = 3. Ordinary one-way ANOVA showed no significant difference between DMSO and different KH-103 concentrations (F(3, 8) = 1.82, p = 0.2215). Follow-up Dunnett’s multiple comparisons tests also showed no significant differences. **e** Immunoblot and quantification of MPro fused with FKBP12^F36V^ transiently expressed in A549 cells treated with KH-103 and dTAG13. For both MPro-N and −C, N = 3 for DMSO, and N = 4 for KH-103 and dTAG13 treatments. Statistical details for the MPro (N) and (C) are summarized in supplementary table 4. P-values # < 0.1, ** < 0.01.

The DEX up-regulated genes in cluster 3 (Fig. 8a) were also up-regulated by MIF and by CORT113176 treatments alone. These genes showed higher GR signals near their TSS compared to the other clusters, suggesting that paradoxically, the inhibitors partially activate GR and showed agonistic activity. The DEX down-regulated genes in cluster 1, were also down-regulated by MIF and CORT113176 but not by KH-103. These genes showed less GR signal near TSS, or enhancer compared to the up-regulated genes upon DEX (cluster 3), indicating indirect regulation or consequence of the transcriptional machinery being recruited to other sites. The DEX-insensitive genes of cluster 2 and cluster 4 were up-regulated and down-regulated, respectively, by the inhibitors and did not show a clear TSS GR binding. This additionally points towards the indirect or off-target activity of the inhibitors (Fig. 8a). Instead, most of the 13 genes triggered by KH-103 have a GR peak in their TSS or putative enhancer (Fig. 8b), potentially indicating some KH-103-mediated activation, but almost no off-target effects.

Interestingly, cytochromes P450 (Cyps) genes were triggered by DEX, MIF, and CORT113176 but not by KH-103 (supplementary Fig. 11). These genes are triggered as a response to initiate xenobiotics metabolism and are known to be regulated by the transcription factor Pregnane X receptor (PXR). This further highlights the specificity of KH-103. We verified that only 2 of the 13 KH-103 triggered genes have predicted PXR binding motifs within their TSS while also containing GR binding motifs, arguing against KH-103 triggering PXR-mediated transcriptional regulation of these genes (Fig. 8b)^38^. Overall, KH-103 shows negligible agonism or inverse agonism.

We further tested the specificity of KH-103 at the protein level using immunoblotting. We selected three candidates based on structural similarity (mineralocorticoid receptor (MR)) and in silico putative binding partners for DEX (interleukin-6 (IL-6)) and SARS-Cov2 main protease (MPro))^39^. KH-103 did not induce degradation of MR at multiple concentrations tested (Fig. 8c). Since IL-6, a secreted extracellular cytokine, and the viral enzyme, MPro, are not endogenously present in our cells, a dTAG-containing version of these proteins was transiently expressed in HEK293. Both IL-6 and MPro were depleted by the control dTAG13 PROTAC, showing that these proteins are, in principle, degradable by the PROTAC approach. Importantly, KH-103 did not induce the degradation of these proteins. Altogether these findings further support the specific targeting of GR by KH-103 (Fig. 8d, e).

### KH-103 allows interrogating the role of GR in regulating neuronal activity in primary culture

Primary cultures constitute the preferred option to study complex neuronal circuit dynamics, which are based on intercellular connectivity. Such properties cannot be interrogated in immortalized cell lines that are amenable to a genetic toolset^40^. Hence, clean pharmacological manipulations are of prime importance. To demonstrate the applicability of KH-103 in primary culture systems, we used calcium imaging in rat cortical primary neurons. It is known that GCs can shift the Ca^2+^ baseline in the rat primary culture with functional consequences on excitability^41^. The effects of prolonged GCs on spontaneous firing in primary rat cortical culture are unknown, and so is the dependency of such effect on GR.

We recorded calcium signals for 1 min at 10 Hz in four different treatment conditions. These included cultures pretreated on the first day with either KH-103 or vehicle and were subsequently supplemented with either DEX or vehicle on the second day (Fig. 9a). Measuring the spontaneous peak-to-peak intervals of the calcium signal from individual neurons, we observed that prolonged DEX treatment increased the intervals of spontaneous Ca^2+^ peaks (Fig. 9b). Importantly, KH-103 alone did not influence the intervals, indicating that the mere presence of inactive GR, e.g., as a scaffold or binding partner for other proteins is irrelevant. Pretreatment with KH-103 effectively blocked the DEX-induced increase in intervals. This suggests a GR-dependent decrease in spontaneous activity in response to DEX and highlights KH-103 as an effective tool for studying the role of GR in primary culture.

**Figure. 9:**
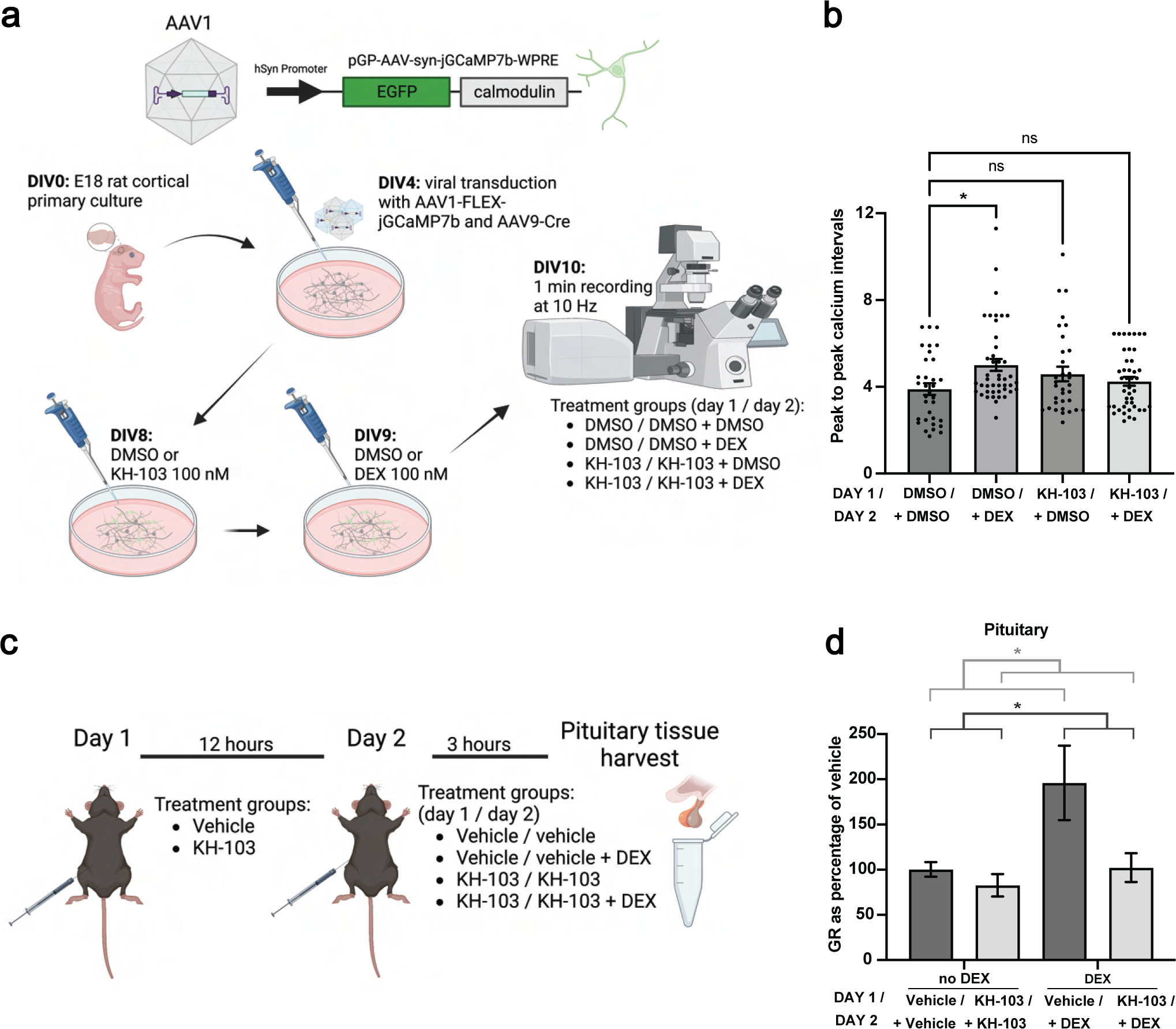
KH-103 application in primary neurons and in-vivo stress mouse model. **a** Schematic depiction of jGCamP7b calcium sensor cassette in AAV viral construct as well as experimental timeline and treatment groups in rat primary cortical neurons. E18: embryonic day 18, DIV: days in-vitro. **b** Peak-to-peak intervals of calcium signals. Each data point (N) is the average of all intervals for each neuron during 60 s recording. N = 33 for DMSO and KH-103, N = 44 for DEX, N = 43 for DEX + KH-103. Two-way ANOVA showed a significant interaction between KH-103 and DEX (F(1,149) = 7.17, p = 0.0082). Follow-up Dunnett’s multiple comparison tests showed a significant difference between DMSO and DEX (p = 0.0113). **c** Schematic depiction and timeline of KH-103 and DEX delivery and pituitary sample collection in male mice. **d** GR levels in the pituitary as assessed by immunoblotting (each group, N = 4). Two-way ANOVA showed a significant effect of DEX (F(1,12) = 6.13, p = 0.0292) and KH-103 (F(1,12) = 5.71, p = 0.0341). P-values * < 0.05.

### In-vivo injection of KH-103 counteracts the effects of DEX on GR in the pituitary

Lastly, since the GR-PROTAC, KH-103 showed robust GR degradation in-vitro, we aimed to assess its degradation capabilities in-vivo in mice. A prime area of high GR levels and crucial for negative feedback inhibition on the HPA axis is the pituitary. We thus evaluated whether KH-103 could effectively interfere with GR activation via DEX in the pituitary (Fig. 9c). To this end, we systemically administered DEX or DEX in combination with KH-103 and assayed GR levels using immunoblotting. We observed a significant rise in GR upon DEX-only injection, an effect that was reversed in the group receiving a prior KH-103-only injection followed by a mixed KH-103/DEX intraperitoneal injection (Fig. 9d). This indicates that while GR levels increase significantly upon GR activation in the pituitary, this effect can be counteracted with KH-103.

## Discussion

We developed a new potentially translatable tool to abolish GR functions at the protein level. Our extensive characterization demonstrates very high efficiency of depletion that is comparable to known PRTOACs against other protein targets, tested in HEK293 cells and even to PROTACs now under evaluation for clinical application directed against the androgen receptor^42,43^.

KH-103 is highly specific, showing no marked cross depletion of the other tested proteins known to have the binding potential for DEX^39^. In the absence of GR activation, KH-103 further shows neither notable agonistic nor inverse agonistic activity. In fact, activation by KH-103 seems unlikely due to the size difference between KH-103 and DEX. Should KH-103 engage in the formation of a tertiary complex, it would likely prevent the complex from binding or forming a heterodimer with other transcription factors. Although based on the candidate-based RT-qPCR experimental approach, we cannot exclude any fast-acting genome-wide agonistic effects, our sequencing results corroborate such interpretation. We hypothesize that this absent agonism is achieved by a complete loss of GR binding to DNA, whereas MIF still retains this ability. The same holds true for selective agonists/antagonists, which are believed to gain their selectivity by modulating specific cofactor complex formation with GR, a process that largely contributes to the context-dependent gene expression regulation via GR^44^.

Jointly with the absence of agonistic activity, the high specificity contributes to the outstanding passive blockage of DEX-induced gene expression changes that we observe in A495 cells (Fig. 7). In the employed tests here, KH-103 shows enhanced blockage of DEX-induced gene expression in comparison to the only clinically approved inhibitor MIF. The versatile applicability in various cellular systems and species is evidence of great potential for future use in and beyond basic research.

In a prior study, a PROTAC was developed against tubulin-associated unit (TAU) protein and showed poor blood-brain-barrier (BBB) penetration efficacy^45^. Hence, to avoid the impact of potential low brain penetration, we chose to assess the efficacy of depleting GR in the HPA axis outside the BBB in the pituitary. The DEX-triggered increase in GR protein is in line with the role of GR in the anterior pituitary in negative feedback inhibition to terminate the stress response, where activation of GR triggers GR homodimerization and induces genomic effects leading to a transient rise in GR levels^46^. In our proof of principle experiment, we show that KH-103 efficiently depletes GR in-vivo, independent of GR activation. Thus KH-103 has the potential to modulate the negative feedback inhibition since it can effectively deplete GR protein, independent of activation. This independence is particularly interesting considering the circadian fluctuations of GC levels, where GR is mostly inactive at daily baseline levels, rendering activation-dependent approaches difficult. At the same time, the possibility of achieving depletion despite high resynthesis rates upon activation in the pituitary emphasizes the high potency of KH-103, making us optimistic about extrapolating toward a general in-vivo efficacy. Future in-vivo experiments could address BBB permeability and dose dependency.

In terms of strategies for novel technology development, we demonstrate that the design of new compounds can harness known ligands of proteins of interest to assemble them into PROTAC compounds, including the possibility to turn known agonists, such as DEX, into an “antagonist”. This strategy can be particularly helpful when the inhibition of a protein-catalytic function is not sufficient, but complete depletion is desirable. The PROTAC-mediated protein depletion approach may - in the future - also prove useful to target other steroid receptors that are involved in all sorts of pathologies.

Overall, we present our GR-PROTAC as a promising, translatable tool to achieve passive abolishment of GR’s genomic function and non-genomic functions. GR-PROTAC complements the available toolbox to study GR in the context of health and disease, an endeavor that has to date, been difficult due to the cross-reactivity of full inhibitors and partial agonistic actions of many selective inhibitors. Besides benefits for basic research on steroid-mediated cellular and stress axis function, KH-103 holds great potential for clinical application in the context of stress-related neuropsychiatric disease, Cushing’s disease but also for the treatment of GR-associated cancer metastasis.

## Materials and Methods

### GR-PROTACs design and synthesis

Synthetic details on all PROTACs are provided in the supplementary Method file and supplementary Fig. 1 and Fig. 2.

### In-vitro cell culture

HEK293 and N2a cells were maintained in Dulbecco’s Modified Eagle Medium with 10% Fetal Bovine serum (FBS) and 1% Penicillin/Streptomycin (P/S) (Gibco, Invitrogen). A549 cells (CCL-185) were purchased from American Type Culture Collection (ATCC) and were maintained in an F-12 medium with 10% FBS and 1% P/S. All cells were kept at 5% CO_2_ at 37 °C.

### Treatment of cells with compounds

KH-95, KH-99, KH-102, KH-103, and DEX (Sigma) were used at 100 nM except indicated otherwise. MIF (Sigma-Aldrich) and CORT113176 (courtesy of Corcept) were used at 1 μM. MG-132 (MedChemExpress) was used at 20 μM. All mentioned compounds were dissolved in DMSO.

### Protein extraction and immunoblotting

Cells were lysed in RIPA buffer (Invitrogen) containing 1x Protease inhibitor (Roche) by incubating on ice for 20 min followed by 10 min centrifugation at 13 k rpm to separate protein lysate from the cell debris. Protein lysates were resolved on 10% gels (mini-PROTEAN TGX, Bio-Rad). For the fractionation experiment, the nuclear fraction proteins were resolved on gradient 8-16% gels (mini-PROTEAN TGX, Bio-Rad). Resolved proteins were transferred to nitrocellulose membranes (Trans-blot Turbo, Bio-Rad) on semi-dry Bio-Rad systems. Blots were incubated for 1 h in 5% milk-TBST and overnight at 4 °C with the primary antibody. After washes, blots were incubated with secondary antibodies for 1 h and were consequently developed using Clarity Western ECL and Clarity Max Western ECL substrates (Bio-Rad). Protein bands were visualized on the ChemidocTM MP imaging system (Bio-Rad).

### Antibodies

The following antibodies were used: GR (G-5, Santacruz sc-393232, 1:100), MR (clone 6G1, Merck MABS496, 1:1000), alpha-tubulin (11H10, cell signaling 2125S, dilution 1:1000), GAPDH (ABS16, Merck Millipore, 1:1000), HA (C29F4, cell signaling #3724, 1:1000), Calpain (Abcam ab28258, 1:1000), VDAC1 (Abcam ab15895, 1:1000), IL-6 (Abcam ab259341), H3 (Abcam ab1791, 1:1000), and GFP (Abcam, ab290, 1:1000). Secondary antibodies included: goat anti-mouse IgG antibody (Merck Millipore AP308P, (H+L) HRP conjugate, 1:20’000) and goat anti-rabbit IgG (Merck Millipore AP307P, (H+L) HRP conjugate, 1:20’000).

### Immunofluorescence staining

Cells were seeded and treated on 0.1% gelatin-coated coverslips. After a wash with 1x PBS, they were fixed in 4% PFA for 20 min at RT, followed by 3x washes in PBS before permeabilization in 0.3% Triton in PBS for 30 min at RT. After 3x further wash steps, cells were blocked overnight at 4 °C in 1% bovine serum albumin (BSA). Primary GR antibody (G-5, Santacruz sc-393232, 1:100) or MAP2 antibody (Invitrogen, 1:2000) incubations lasted for 1 h at RT followed by 3x washes and 1 h incubation in secondary antibody at RT (Cy3 Goat anti-mouse against GR, Jackson ImmunoResearch 115-165-003, 1:300 and goat anti-chicken Alexa488 against MAP2, Thermo Fisher Scientific A-11039, 1:1000). After 3x washes, coverslips were mounted on glass slides using fluoroshield mounting medium plus DAPI fluorescent nuclear stain (Abcam). Slides were dried, sealed, and stored at 4 °C until imaging.

### Image acquisition – Confocal/fluorescence microscopy

All immunofluorescence images were acquired using a Zeiss LSM 880 confocal microscope or Zeiss Axio observer 7 wide field inverted microscope at a specified magnification. Images were taken with ZEN 2.3 Pro software. All laser intensities and gamma values always remained the same across all conditions. Scale bars were added, and adjustments were made in ZEN 2.6 (Blue) software.

### Fractionation

Cells were seeded in 100 mm dishes. The Fractionation protocol was followed according to Baghirova S. et al., MethodsX, 2015 to acquire organelle membrane, cytosolic and nuclear fractions^47^. In brief, cells were collected by scraping and were centrifuged at 500 xg at 4 °C for 10 min. Pellets were washed with 1x PBS and were centrifuged again. This time pellet was lysed in ice-cold buffer A (NaCl 150 mM, HEPES (pH 7.4) 50 mM, Digitonin (Sigma, D141) 25 ug/ml, Hexylene glycol (Sigma, 112100) 1M) containing protease inhibitor (Roche). After 10 min incubation on an end-over-end rotator at 4 °C, the lysate was centrifuged at 2000 xg for 10 min at 4 °C. Supernatant was collected and stored at 80 °C until further analysis as the cytosolic fraction. The pellet was lysed in ice-cold buffer B (NaCl (150 mM), HEPES (pH 7.4) 50 mM, Igepal (Sigma, I8896) 1% v:v, Hexylene glycol (Sigma, 112100) 1M containing protease inhibitor (Roche). Lysate was incubated on ice for 30 min and subsequently centrifuged at 7000 xg for 10 min at 4 °C. Supernatant was collected as the membrane-bound organelle protein fraction (except those from the nucleus) and was stored at 80 °C until further analysis. Pellet was lysed in ice-cold buffer C (NaCl 150 mM, HEPES (pH 7.4) 50 mM, Sodium dodecyl sulfate (Carl Roth, CN30.3) 0.1% w:v, Hexylene glycol (Sigma, 112100) 1M containing Benzonase (Sigma, E1014) and protease inhibitor (Roche) and was incubated on an end-over-end rotator for 30 min at 4 °C. This lysate was further sonicated for 3x 10 s in a cold-water bath before centrifugation at 7800 xg for 10 min at 4 °C. Supernatant contained the nuclear protein fraction, which was stored at 80 °C until the immunoblotting analysis.

### RNA extraction

Total RNA was extracted using Quick-RNA Microprep Kit (Zymoresearch, R1051), followed by cDNA synthesis. RNA concentration was measured by NanoDrop, and integrity was checked on an agarose gel.

### C-DNA synthesis

500 ng of total RNA were converted to cDNA using M-MLV Reverse Transcriptase (Promega, M1705) and Oligo(dt) 15 primers (Promega, C1101) containing RNasin® Plus RNase inhibitor (Promega, N2615) according to the manufacturer’s instructions.

### RT-qPCR

cDNAs were diluted 1:5, and 2 μl of that was assessed in RT-qPCR in triplicates. HPRT and PPIA housekeeping genes were used to generate normalization factors using the geNorm algorithm^48^. Sybr green Mix (Roche) was used for the amplification signal, and the plates were run on a CFX384 Real-Time PCR system (Bio-Rad) device. Primers can be found in the supplementary source data.

### RNA sequencing library preparation and sequencing

For the gene expression studies, A549 cells were treated with DMSO, KH-103, DEX, MIF, CORT113176, or combinations for varying times (please see supplementary Fig. 3 for the immunofluorescent staining of the treatment conditions). Total RNA was isolated using the Quick-RNA Microprep Kit (Zymoresearch, R1051). RNA samples at 25 ng/μl concentration were processed at Novogene, UK.

Non-directional Poly-A library preparations were performed according to the manufacturer’s recommendations (Next® Ultra RNA Library Prep Kit for Illumina®). Briefly, mRNA was purified using poly-T oligo-attached magnetic beads, followed by fragmentation and first-strand cDNA synthesis using random hexamer primers. The second strand of cDNA was then synthesized using dTTP. This was followed by end repair, A-tailing, adapter ligation, and size selection. Next, libraries were PCR amplified and purified. For quality control and quantification, libraries were checked with Qubit and RT-qPCR. Size distribution was assessed on a bioanalyzer (Agilent 2100). Quantified libraries were then pooled. Clusters of index-coded samples were generated, and libraries were sequenced on Illumina NovaSeq 6000 S4 flowcell with PE150 to generate 6G of paired-end reads.

### Sequencing data analysis

Gene expression was quantified from RNA-seq data using Salmon 1.7.0 with the –validateMappings option on the Ensembl 105 transcriptome^49^. Counts were aggregated to gene level using the tximport package^50^. For differential expression analysis, all samples were considered together, and two surrogate variables were included in the model using the sva 3.44.0 R package^51^. Differential expression analysis was performed with edgeR 3.38.1, filtering genes using filterByExpr with a minimum count of 20^52^. For the publicly available ENCODE data, we obtained processed data from the encodeproject.org website (see GitHub repository for exact files). For RNA-seq, we used gene count tables, and for chromatin data, we used the signal p-value bigwig and the IDR-thresholded peaks across replicates. Signal heatmaps were generated with the epiwraps R package 0.99.50. To select the ’best putative distal enhancer’ for given genes, we used enhancer target predictions and selected the predicted enhancer with the highest GR signal^53^.

### Plasmid constructions

Codon-optimized cDNA of GR isoforms as well as MPro were synthesized by IDT and were cloned into entry plasmid Gateway™ pDONR™221 Vector (Thermo Fischer) before using LR reaction to transfer the cDNA into destination vectors pLEX_305-C-dTAG or pLEX_305-N-dTAG (Addgene #91797 & #91798, gift from James Bradner & Behnam Nabet) using gateway cloning method (Thermo Fisher)^54^. For cDNA sequences, gateway, and sequencing primers, please see supplementary source data. Plasmid pEGFP-GR (Addgene #47504, gift from Alice Wong) was used to express GR-EGFP, and pEGFP-N1-IL6 (Addgene #111933, gift from Geert van den Bogaart) plasmid was used for expression of IL-6-EGFP^55^. The absence of mutations in the inserted cDNAs was verified prior to use.

### Neuronal primary culture

Hippocampi or cortices of E18 pups were isolated and dissociated in 37 °C TryPLE for 7 min with inverting. TryPLE was then removed by twice washing the digested tissue in the dissection medium (Leibovitz’s L-15 Medium (Gibco, 11415-04) + 7 mM HEPES (Milian, L0180-500)). Tissues were then fully dissociated by applying mechanical force using a pipette in warm Neurobasal plus (NB+) medium (Thermo Fisher, A3582901) supplemented with 2 mM GlutaMAX, 2% B27, 1% P/S (Gibco, Invitrogen). Neurons were then counted and seeded on PLL-coated coverslips and maintained in NB+ medium in 5% CO_2_ at 37 °C. Half of the media was refreshed every third day.

### Calcium imaging and analysis

E-18 rat primary cortical neurons were cultured on polyethyleneimine (Sigma-Aldrich) and laminin (Sigma-Aldrich) coated coverslips. Cells were transduced with floxed jGCaMP7b (adeno-associated viruses (AAV) 1-syn-FLEX-jGCaMP7b-WPRE; Addgene #104493, MOI = 5 × 10^5^ vg, a gift from Douglas Kim & GENIE Project) and the same titer of Cre (AAV9-hSyn-Cre-WPRE-hGH; Addgene #105553, MOI = 5 × 10^5^ vg, a gift from James M. Wilson) AAVs at days in-vitro (DIV) 4^56^. Cells were treated with compounds on DIV 8 and 9. Calcium signals were recorded on DIV 10 using a Nikon NiE upright microscope equipped with Yokogawa W1 spinning disk scan head for 1 min at 10 Hz. Calcium traces were extracted using the software Suite2p^57^.

### Animals

Animals were housed in individually ventilated cages on an inverted 12 h light-dark cycle (lights on at 08:15 am). Food and water were provided ad libitum. They were derived from in-house breeding colonies and acclimated for at least 1 week to the respective housing room. All experiments were conducted complying with ethical regulations and local animal guidelines (according to Swiss accreditation License number: ZH067/2022).

### Injections

To prepare the stock, KH-103 powder was solved in 40% Captisol (Selleck Chemicals) in DMSO. To prepare the injection solutions, 5% of the KH-103 in Captisol/DMSO was formulated in 20% Solutol (GLPBIO) in 0.9% sterile saline (Moltox) (w:v) solution^33,58^. The final formulation contained 5% DMSO. The mixture was well vortexed and sonicated in a heated water bath until achieving a homogeneous solution ready for injection.

### Statistics

All statistics, except for the RNA sequencing data analysis, were performed using the built-in statistical function in GraphPad Prism. The confidence level was consistently set to 95%, and hence p-value below 0.05 was considered statistically significant. All error bars are represented as the standard error of the mean. The used test and statistical parameters are specified in the figure captions or in the mentioned supplementary tables.

## Data availability

The code underlying the bioinformatic analysis, including processed data objects, are available on the GitHub repository: https://github.com/ETHZ-INS/Glucorticoid-protacs. GR-Chip-Sequencing data were accessed from the T. Reddy lab publication^6^. Sequencing data have been deposited at GEO; the accession number is available upon request. The remaining data are being deposited to ETH research collection and can also be made available upon request.

## Author contributions

K.M.H., under the supervision of E.M.C., designed and synthesized GR-PROTAC compounds, prepared illustrations, and helped with results interpretation. E.M.C provided resources for PROTAC synthesis. M.G., with the help of S.M.B., K.M., and R.R., performed all the characterizations in immortal cells (immunoblotting and IFS experiments, ligand competition experiments, fractionation, GR isoform experiments, and RT-qPCR experiments). M.G. performed the mouse primary neuronal cultures, the IL6 and MPro experiments, RNAseq treatments, and RNA extraction. P.L.G and D.P. performed the RNA seq analyses and generated graphs. M.G. and X.X., under the supervision of A.H., performed the calcium imaging experiment. M.G., M.P., M.K., and V.F. performed mouse studies with help from I.I.. M.G. prepared illustrations and graphs. M.G. and K.G. designed the experiments, interpreted the results, and wrote the manuscript with inputs from K.M.H., P.L.G., O.M., and J.B.. K.G. provided resources.

## Acknowledgments

We thank Hazel Hunt at Corcept for constructive comments and for providing CORT113176 and Behnam Nabet for his advice on in-vivo drug delivery formulations and for providing the dTAG constructs. The Gapp lab receives funding from an SNF PR00P3_201543, an SBFI-funded ERC starting grant REF-1131-52105, an ETH Project Grant ETH-41 20-1, and the Olga Mayenfisch foundation. Mahshid Gazorpak received funding from the Swiss Hirnliga. Vincent Fischer is funded by a ZNZ-Ph.D. fellowship. The Bohacek lab was supported by the ETH Zurich, ETH Project Grant ETH-20 19-1, the Swiss National Science Foundation (grants 310030_172889/1 and 310030_204372), the Swiss 3R competence center and the Botnar Research Centre for Child Health, Multi-Investigator Project. The Hierlemann lab receives funding from the ERC Advanced Grant 694829 ‘neuroXscales’. Illustrations were made by BioRender.

## List of abbreviations

AAV: Adeno-associated virus
ALS: Amyotrophic Lateral Sclerosis
BBB: Blood-brain-barrier
Bp: Base pair
ChIP-seq: Chromatin immunoprecipitation sequencing
CPM: Counts per million
CRBN: Cereblon
Cyps: Cytochromes P450
DEG: Differentially expressed gene
DEX: Dexamethasone
DMSO: Dimethyl sulfoxide
dTAG: Degradation TAG
Dusp1: Dual-specificity phosphatase 1
EGFP: Enhanced green fluorescent protein
FDR: False discovery rate
FKBP12: FK506-binding protein 12
FKBP5: FK506-binding Protein 51
GC: Glucocorticoid
GR: Glucocorticoid receptor
H: Hour(s)
HEK293: Human embryonic kidney 293
Histone 3: H3
HPA: Hypothalamus-Pituitary-Adrenal
Hz: Hertz
IL-6: Interleukin-6
KD: Dissociation constant
Kda: Kilodalton
LBD: Ligand binding domain
logFC: Log fold change
MAP2: Microtubule-associated protein 2
MIF: Mifepristone
Min: Minute(s)
MPro: SARS-CoV-2 main protease
MR: Mineralocorticoid receptor
N2a: Neuro 2a
PDB: Protein data bank
PEG: Poly(ethylene glycol)
Per1: Period circadian regulator 1
PROTAC: Proteolysis targeting chimeras
PXR: Pregnane X receptor
RT-qPCR: Reverse transcription-quantitative polymerase chain reaction
Sgk1: Serum/glucocorticoid regulated kinase 1
TAU: Tubulin-associated unit
TSS: Transcription start site
UPS: Ubiquitin-proteasome system
VDAC: Voltage-dependent anion channel

