## Supplementary material and tables for "Harnessing PROTAC technology to combat stress hormone receptor activation"

Supplementary Fig. 1-2_Gazorpak et al.

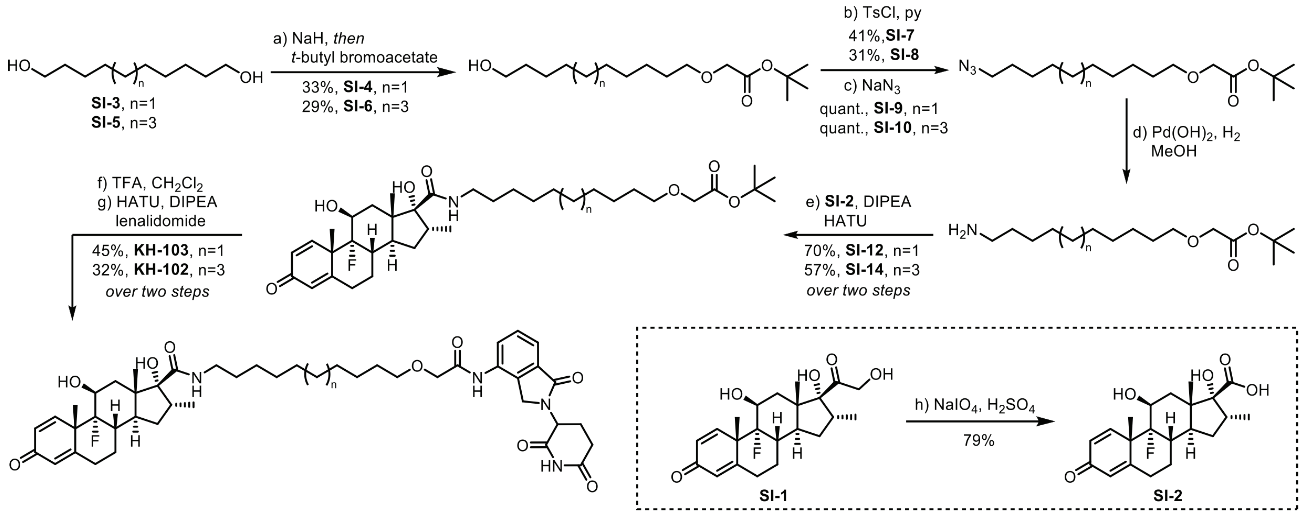


Supplementary Fig. 1_Synthesis of PROTAC KH-102 and PROTAC KH-103. Reagents and conditions: (a) NaH, t-butyl bromoacetate, r.t., 16 h, 33% for SI-4, 29% for SI-6; (b) TsCl, pyridine, 0 °C to r.t., 2 h, 41% for SI-7, 31% for SI-8; (c) NaN3, DMF, 100 °C, 3 h, quant. for SI-9, quant for SI-10; (d) Pd(OH)2, H2, r.t., 1.5 h; (e) SI-2, HATU, DMF, DIPEA, r.t., 16 h, 70% for SI-12, 57% for SI-14; (f) TFA, CH2Cl2, r.t., 1.5 h; (g) lenalidomide, HATU, DMF, DIPEA, r.t., 16 h, 45% for KH-103, 32% for KH-102; (h) NaIO4, H2SO4, EtOH:water (3:1), r.t., 16 h, 79%.

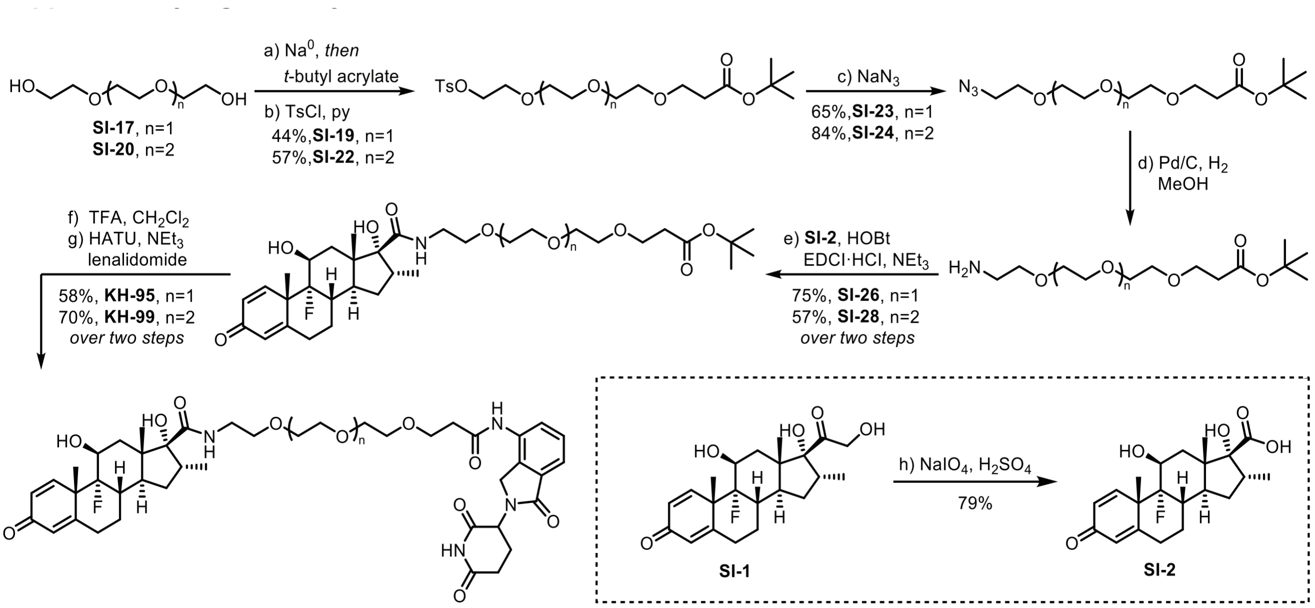


Supplementary Fig. 2_Synthesis of PROTAC KH-95 and PROTAC KH-99. Reagents and conditions: (a) Na0, t-butyl acrylate, r.t., 16 h; (b) TsCl, pyridine, 0 °C to r.t., 16 h, 44% for SI-19, 57% for SI-22; (c) NaN3, DMF, r.t., 16 h, 65% for SI-23, 84% for SI-24; (d) Pd/C, H2, r.t., 16 h; (e) SI-2, EDCI·HCl, HOBt, CH2Cl2, NEt3, r.t., 16 h, 75% for SI-26, 57% for SI-28; (f) TFA, CH2Cl2, r.t., 1.5 h; (g) lenalidomide, HATU, DMF, DIPEA, r.t., 16 h, 58% for KH-95, 70% for KH-99; (h) NaIO4, H2SO4, EtOH:water (3:1), r.t., 16 h, 79%.

Supplementary Fig. 3-4_Gazorpak et al.

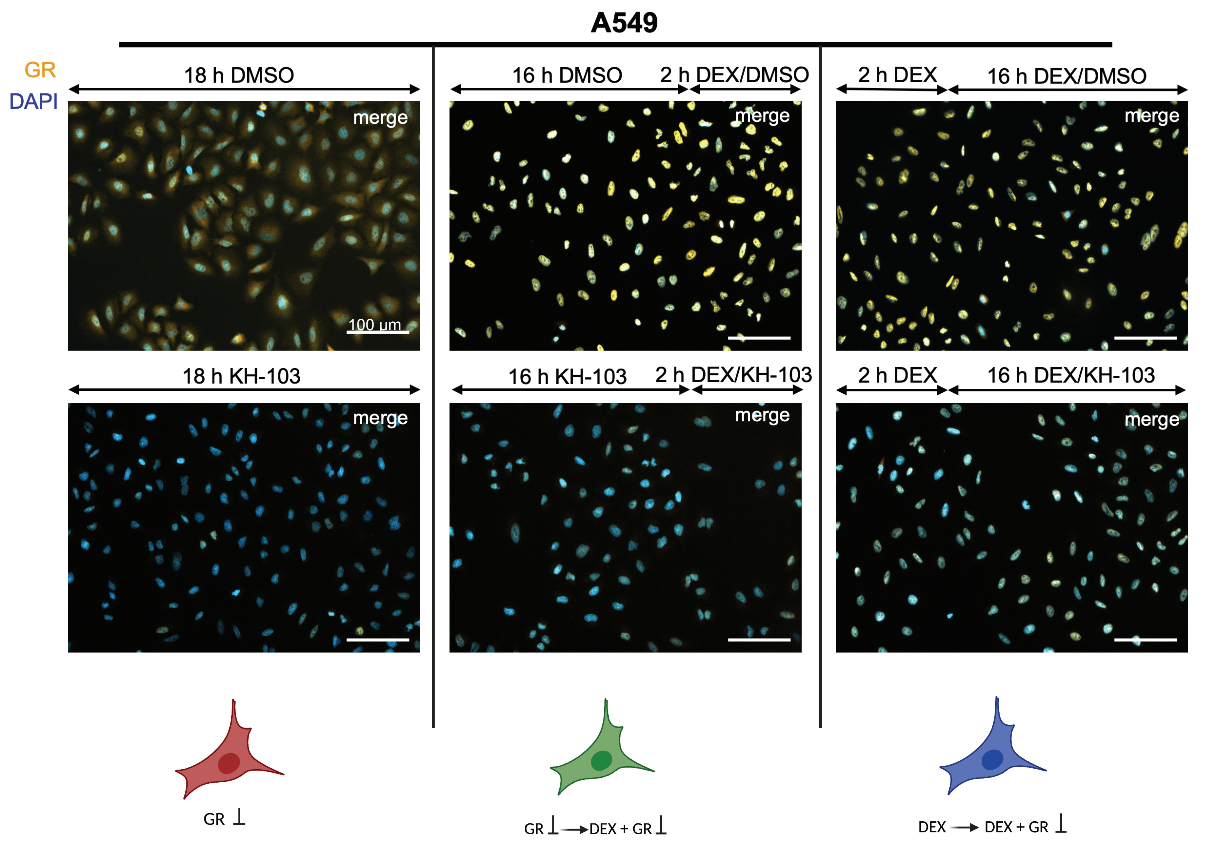


Supplementary Fig. 3_Immunofluorescent staining of GR in A549 cells treated according to the RNAseq experiment conditions described in Fig. 6a. Scale bars: 100 µm

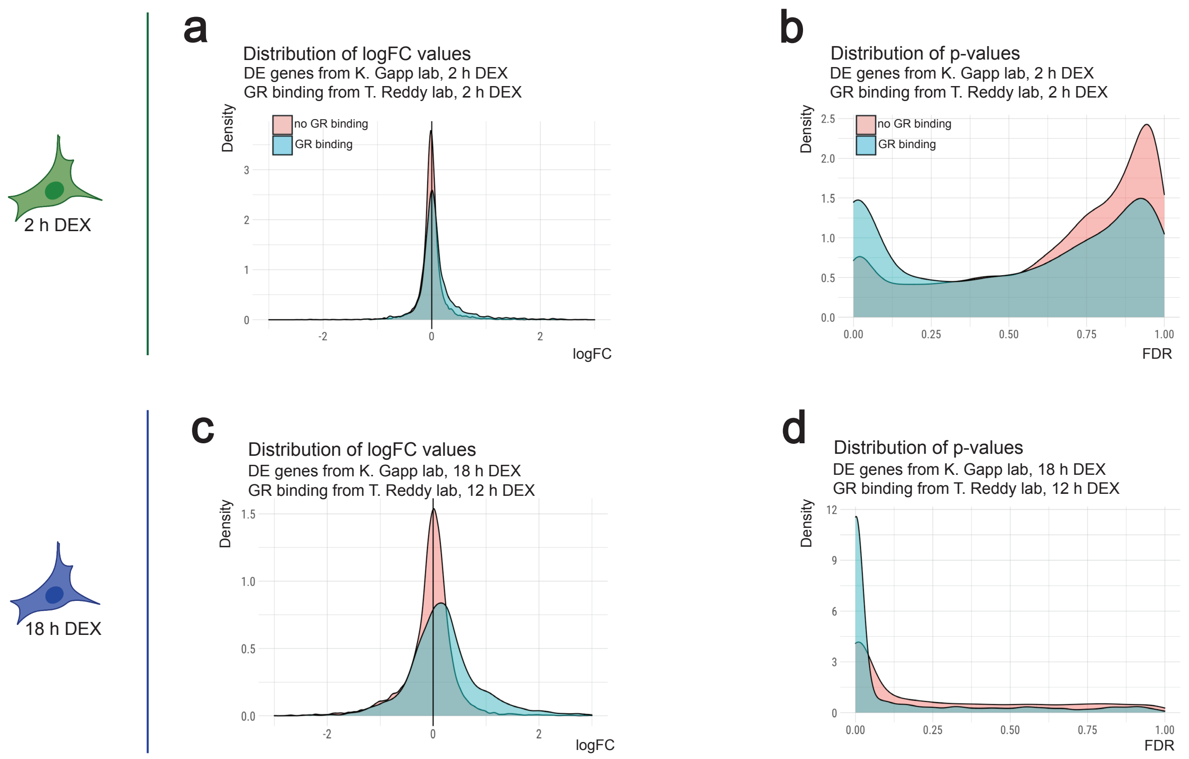


Supplementary Fig. 4_DEGs in 2 h (a) and 18 h DEX treatment (c) with a GR binding signal within + / - 2500 bp from their promoters, showed higher fold changes (shift of the curve to the right or left, away from zero) than those without GR binding signal. This was particularly pronounced at the 18 h time point (c). DEGs in 2 h (b) and 18 h DEX treatment (d) with a GR binding signal within + / - 2500 bp from their promoters, showed higher density of smaller p-values than those without a GR peak.

Supplementary Fig. 5-7_Gazorpak et al.

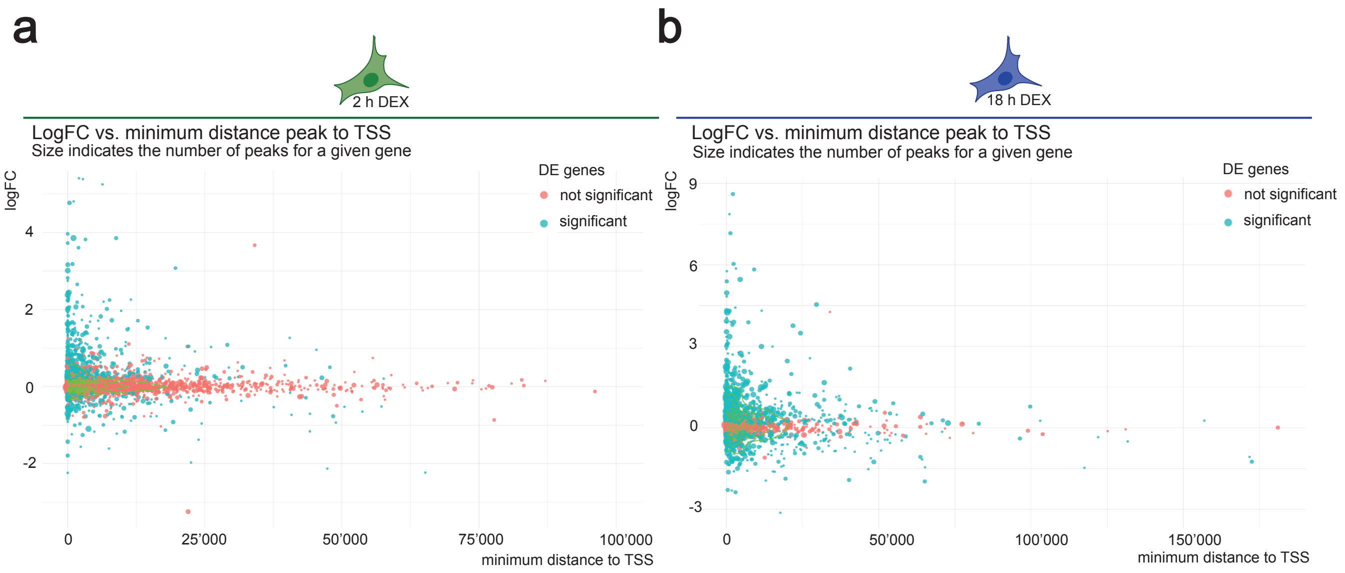


Supplementary Fig. 5_For DEGs in both 2 h (a) and 18 h (b) DEX treatment, the further away the GR binding signal peak from the gene's TSS, the smaller logFC was observed.

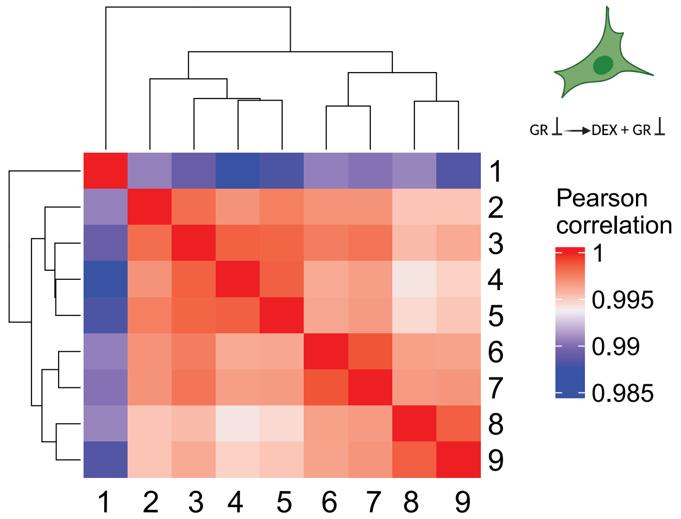


Supplementary Fig. 6_ Pearson correlations and clustering of the conditions based on log(CPM) of the union of DEGs in DEX 2 h condition. 1 : 16 h DMSO > 2 h DEX + DMSO, 2 : 16 h KH-103 > 2 h DEX + KH-103, 3 : 18 h KH-103, 4 : 18 h untreated, 5 : 18 h DMSO, 6 : 16 h MIF > 2 h DEX + MIF, 7 : 18 h MIF, 8 : 16 h CORT113176 > 2 h DEX + CORT113176, 9 : 18 h CORT113176. CPM : Counts Per Million

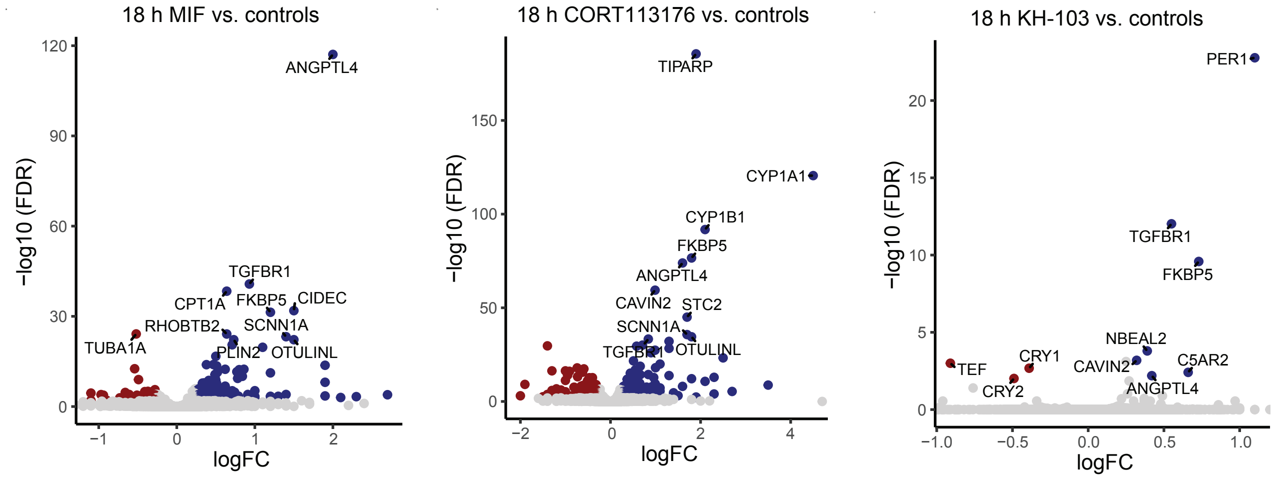


Supplementary Fig. 7_Volcano plots of DEGs in A549 cells treated with MIF (a), CORT113176 (b), and KH-103 (c) for 18 h normalized to controls (untreated and DMSO conditions).

Supplementary Fig. 8_Gazorpak et al.

a

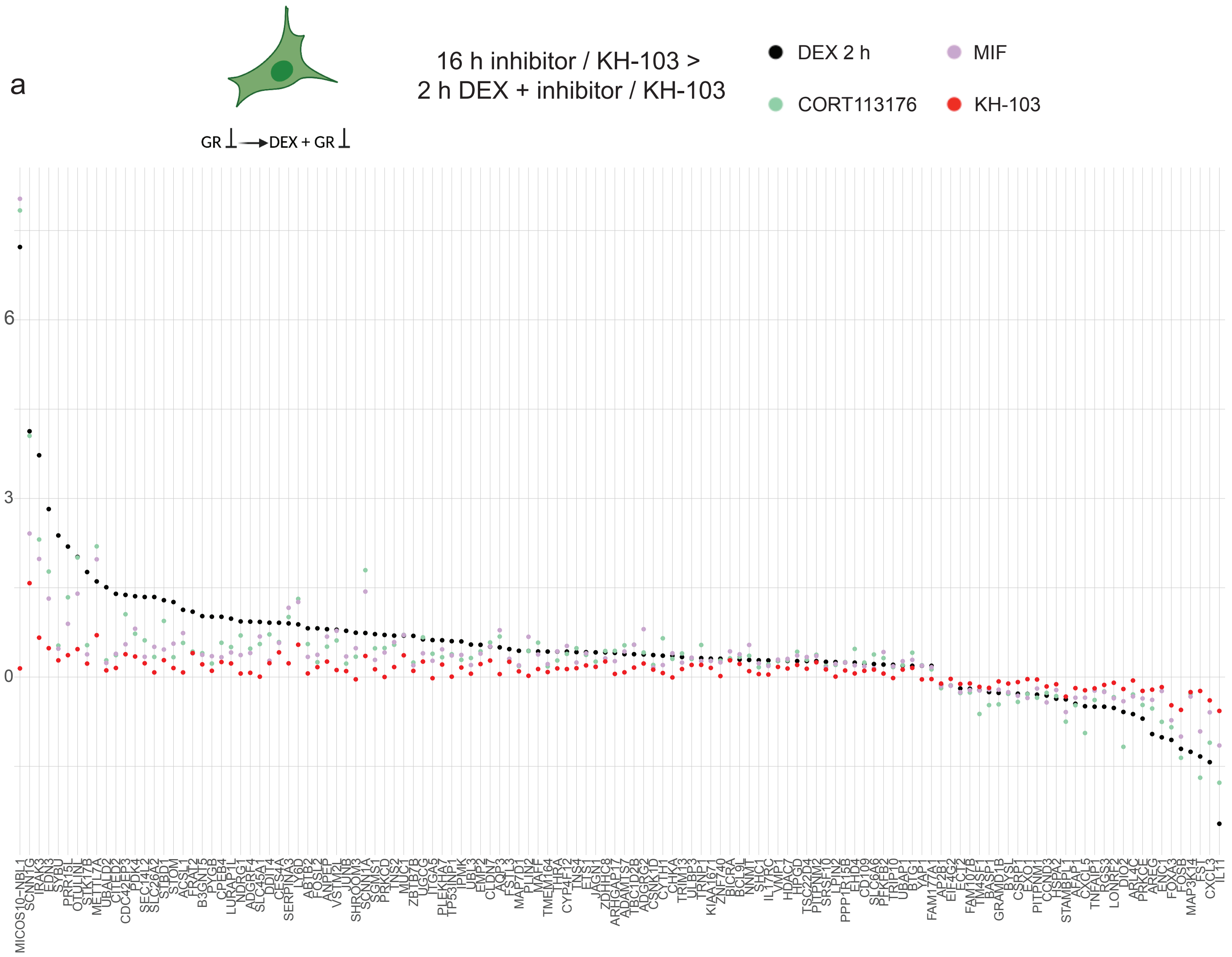


b

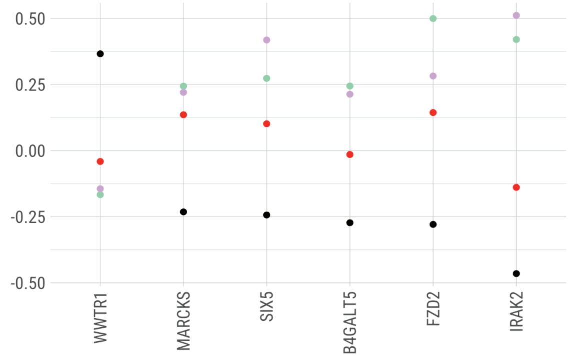


Supplementary Fig. 8_ a genes in the blocking group that showed no significant change upon pretreatment with KH-103 and remained comparable to controls, but they were significantly changed in the same direction as DEX when pretreated with inhibitors (MIF or CORT113176) prior to DEX exposure. b genes that showed no significant change upon pretreatment with KH-103 and remained comparable to controls but were significantly changed upon the inhibitors to the opposite direction as DEX.

Supplementary Fig. 9_Gazorpak et al.

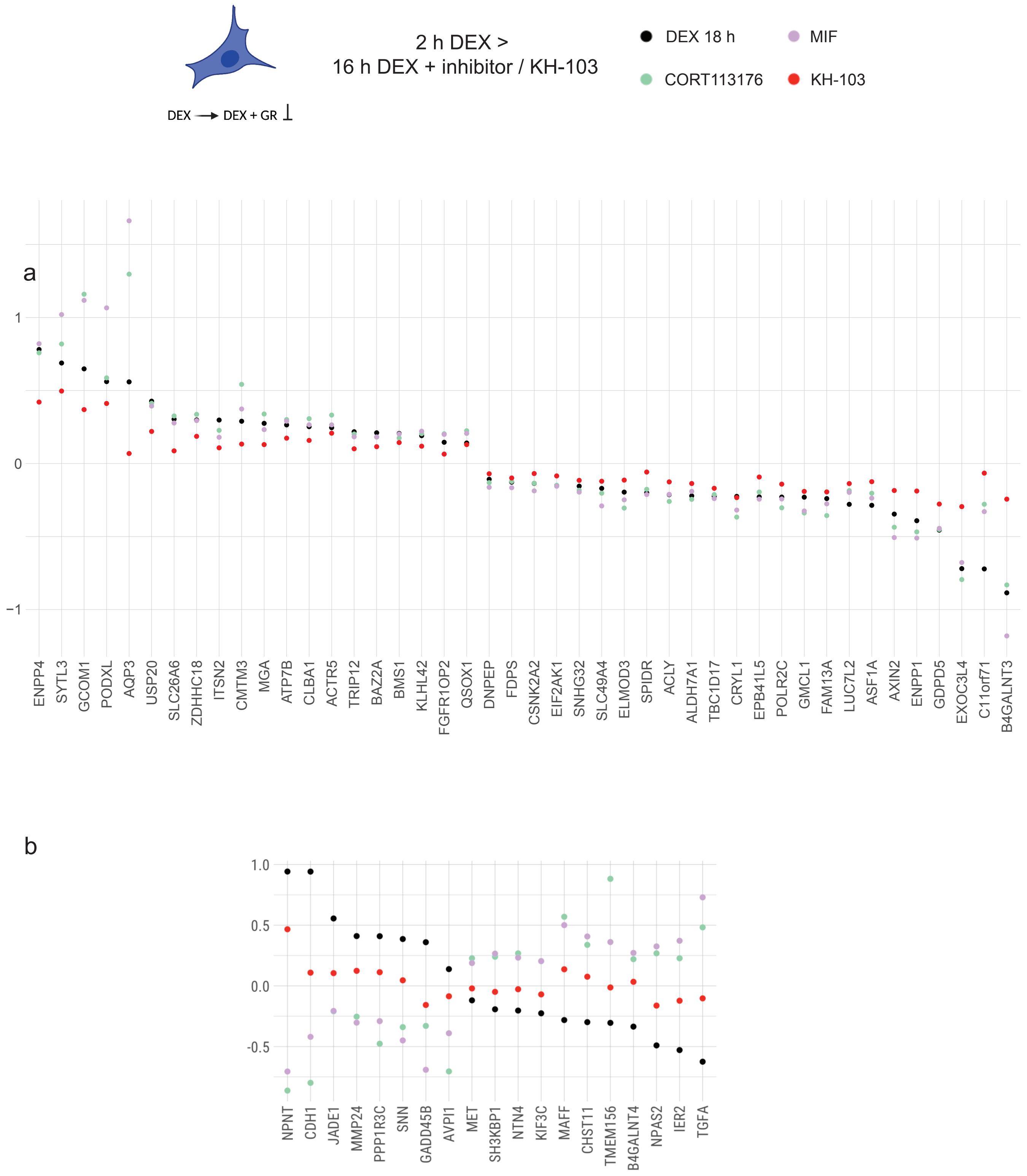


Supplementary Fig. 9_a genes in the reversing group which were significantly changed in the same direction as DEX by the inhibitors, but addition of KH-103 reversed them to comparable levels as controls (were not significantly changed). b genes that showed no significant change compared to controls upon reversing with KH-103 but were significantly changed by the inhibitors in the opposite direction as DEX.

Supplementary Fig. 10-12_Gazorpak et al.

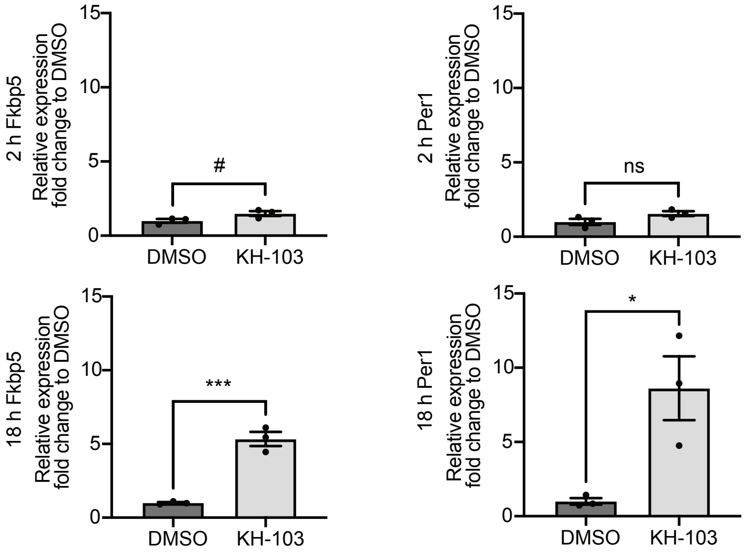


Supplementary Fig. 10_Validation of FKBP5 and Per1 mRNA expression changes upon KH-103 at 2 h and 18 h in A549 cells via qRT-PCR. Results were normalised to HPRT and PPIA housekeeping genes. N = 3. For FKBP5, unpaired, two-tailed t-test showed a trend between DMSO and KH-103 at 2 h ($t(4) = 2.42$, $p = 0.0729$), and a significant difference at 18 h ($t(4) = 8.92$, $p = 0.0009$). For Per1, unpaired, two-tailed t-test showed no significant difference between DMSO and KH-103 at 2 h ($t(4) = 2.10$, $p = 0.1032$), and a significant difference at 18 h ($t(4) = 3.53$, $p = 0.0242$). P-values # < 0.1, * < 0.05, *** < 0.001.

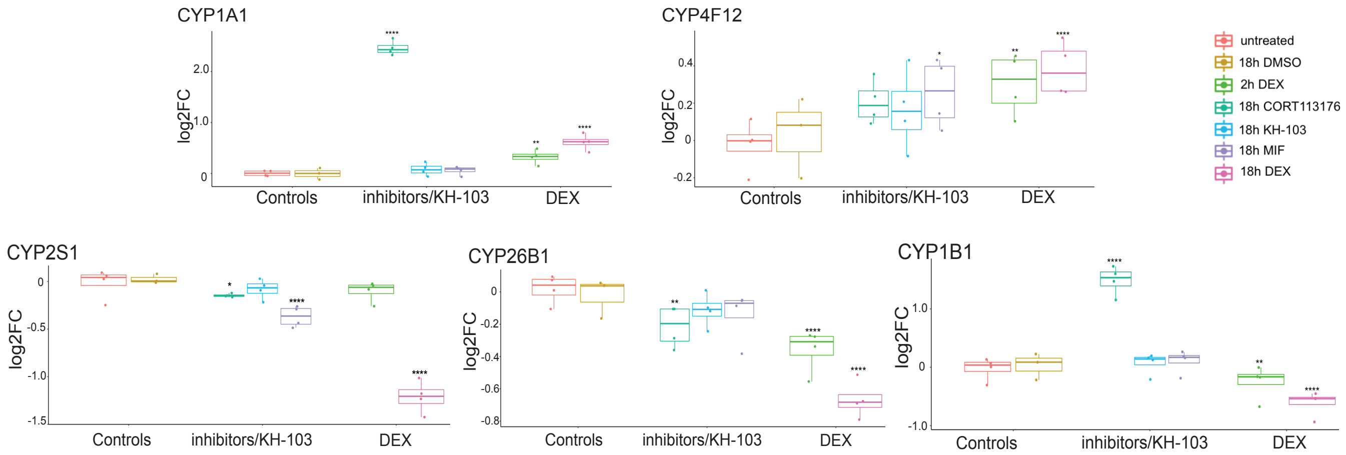


Supplementary Fig. 11_Comparison of CYPs genes expression changes obtained by RNAseq upon 18 h treatment with DEX, or MIF, or CORT113176, or KH-103. Stars represent FDR values # < 0.1, * < 0.05, ** < 0.01, **** < 0.0001, all the logFC and FDR values are summarised in Supplementary Table. 5).

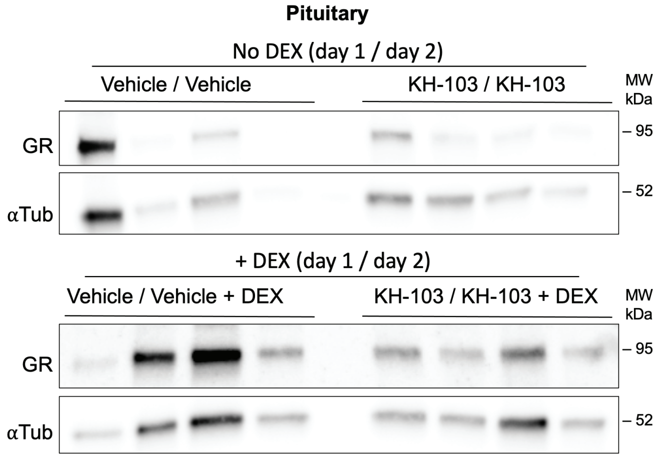


Supplementary Fig. 12_Immunoblot of pituitary tissues in mice injected with vehicle (day 1) / vehicle (day 2), KH-103 / KH-103, vehicle / vehicle + DEX, or KH-103 / KH-103 + DEX (N =4 for each group).

Supplementary Table. 1-5_Gazorpak et al.

| Fisher's LSD | Mean difference | 95% CI of difference | Summary | P-Value |
|---|---|---|---|---|
| A: DMSO/DMSO vs. C: DEX/DEX | 53.67 | 17.87 to 89.46 | ** | 0.0067 |
| A: DMSO/DMSO vs. E: DEX/DMSO | 20.67 | -15.13 to 56.46 | ns | 0.2323 |
| A: DMSO/DMSO vs. B: DMSO/KH-103 | 96.33 | 60.54 to 132.10 | **** | < 0.0001 |
| A: DMSO/DMSO vs. D: DEX/DEX+KH-103 | 62.67 | 26.87 to 98.46 | ** | 0.0025 |
| A: DMSO/DMSO vs. F: DEX/KH-103 | 92.67 | 56.87 to 128.50 | *** | 0.0001 |
| C: DEX/DEX vs. E: DEX/DMSO | -33.00 | -68.79 to 2.79 | # | 0.0676 |
| C: DEX/DEX vs. B: DMSO/KH-103 | 42.67 | 6.88 to 78.46 | * | 0.0233 |
| C: DEX/DEX vs. D: DEX/DEX+KH-103 | 9.00 | -26.79 to 44.79 | ns | 0.5938 |
| C: DEX/DEX vs. F: DEX/KH-103 | 39.00 | 3.21 to 74.79 | * | 0.0351 |
| E: DEX/DMSO vs. B: DMSO/KH-103 | 75.67 | 39.87 to 111.50 | *** | 0.0006 |
| E: DEX/DMSO vs. D: DEX/DEX+KH-103 | 42.00 | 6.21 to 77.79 | * | 0.0252 |
| E: DEX/DMSO vs. F: DEX/KH-103 | 72.00 | 36.21 to 107.80 | *** | 0.0009 |
| B: DMSO/KH-103 vs. D: DEX/DEX+KH-103 | -33.67 | -69.46 to 2.13 | # | 0.0629 |
| B: DMSO/KH-103 vs. F: DEX/KH-103 | -3.67 | -39.46 to 32.13 | ns | 0.8271 |
| D: DEX/DEX+KH-103 vs. F: DEX/KH-103 | 30.00 | -5.79 to 65.79 | # | 0.0928 |

Supplementary Table. 1_Summary of Fisher's LSD multiple comparison of the cytosolic fraction.

| Fisher's LSD | Mean difference | 95% CI of difference | Summary | P-Value |
|---|---|---|---|---|
| A: DMSO/DMSO vs. C: DEX/DEX | 50.67 | 16.84 to 84.49 | ** | 0.0068 |
| A: DMSO/DMSO vs. E: DEX/DMSO | 44.00 | 10.18 to 77.82 | * | 0.0150 |
| A: DMSO/DMSO vs. B: DMSO/KH-103 | 95.67 | 61.84 to 129.50 | **** | < 0.0001 |
| A: DMSO/DMSO vs. D: DEX/DEX+KH-103 | 63.00 | 29.18 to 96.82 | ** | 0.0016 |
| A: DMSO/DMSO vs. F: DEX/KH-103 | 96.00 | 62.18 to 129.80 | **** | < 0.0001 |
| C: DEX/DEX vs. E: DEX/DMSO | -6.67 | -40.49 to 27.16 | ns | 0.6752 |
| C: DEX/DEX vs. B: DMSO/KH-103 | 45.00 | 11.18 to 78.82 | * | 0.0134 |
| C: DEX/DEX vs. D: DEX/DEX+KH-103 | 12.33 | -21.49 to 46.16 | ns | 0.4423 |
| C: DEX/DEX vs. F: DEX/KH-103 | 45.33 | 11.51 to 79.16 | * | 0.0128 |
| E: DEX/DMSO vs. B: DMSO/KH-103 | 51.67 | 17.84 to 85.49 | ** | 0.0060 |
| E: DEX/DMSO vs. D: DEX/DEX+KH-103 | 19.00 | -14.82 to 52.82 | ns | 0.2444 |
| E: DEX/DMSO vs. F: DEX/KH-103 | 52.00 | 18.18 to 85.82 | ** | 0.0058 |
| B: DMSO/KH-103 vs. D: DEX/DEX+KH-103 | -32.67 | -66.49 to 1.16 | # | 0.0571 |
| B: DMSO/KH-103 vs. F: DEX/KH-103 | 0.33 | -33.49 to 34.16 | ns | 0.9832 |
| D: DEX/DEX+KH-103 vs. F: DEX/KH-103 | 33.00 | -0.82 to 66.82 | # | 0.0550 |

Supplementary Table. 2_Summary of Fisher's LSD multiple comparison of the membrane fraction.

| Fisher's LSD | Mean difference | 95% CI of difference | Summary | P-Value |
|---|---|---|---|---|
| A: DMSO/DMSO vs. C: DEX/DEX | -491.00 | -567.10 to -414.90 | **** | < 0.0001 |
| A: DMSO/DMSO vs. E: DEX/DMSO | -234.30 | -310.40 to -158.30 | **** | < 0.0001 |
| A: DMSO/DMSO vs. B: DMSO/KH-103 | 65.00 | -9.68 to 139.70 | # | 0.0822 |
| A: DMSO/DMSO vs. D: DEX/DEX+KH-103 | -228.30 | -304.40 to -152.30 | **** | < 0.0001 |
| A: DMSO/DMSO vs. F: DEX/KH-103 | 64.67 | -11.39 to 140.70 | # | 0.0881 |
| C: DEX/DEX vs. E: DEX/DMSO | 256.70 | 180.60 to 332.70 | **** | < 0.0001 |
| C: DEX/DEX vs. B: DMSO/KH-103 | 571.50 | 486.50 to 656.50 | **** | < 0.0001 |
| C: DEX/DEX vs. D: DEX/DEX+KH-103 | 262.70 | 186.60 to 338.70 | **** | < 0.0001 |
| C: DEX/DEX vs. F: DEX/KH-103 | 555.70 | 479.60 to 631.70 | **** | < 0.0001 |
| E: DEX/DMSO vs. B: DMSO/KH-103 | 314.80 | 229.80 to 399.90 | **** | < 0.0001 |
| E: DEX/DMSO vs. D: DEX/DEX+KH-103 | 6.00 | -70.06 to 82.06 | ns | 0.8653 |
| E: DEX/DMSO vs. F: DEX/KH-103 | 299.00 | 222.90 to 375.10 | **** | < 0.0001 |
| B: DMSO/KH-103 vs. D: DEX/DEX+KH-103 | -308.80 | -393.90 to -223.80 | **** | < 0.0001 |
| B: DMSO/KH-103 vs. F: DEX/KH-103 | -15.83 | -100.90 to 69.20 | ns | 0.6898 |
| D: DEX/DEX+KH-103 vs. F: DEX/KH-103 | 293.00 | 216.90 to 369.10 | **** | < 0.0001 |

Supplementary Table. 3_Summary of Fisher's LSD multiple comparison of the nuclear fraction.

| transcriptional GR isoforms | one-way ANOVA | Holm-Šídák's: DMSO vs. | Summary |
|---|---|---|---|
| GR-α (N) | F(2, 6) = 6.81, p = 0.0286 | KH-103: p = 0.0231 | * |
| | | dTAG13: p = 0.0424 | * |
| GR- γ (N) | F(2, 6) = 13.49, p = 0.0060 | KH-103: p = 0.0292 | * |
| | | dTAG13: p = 0.0041 | ** |
| GR-β (N) | F(2, 6) = 7.82, p = 0.0213 | KH-103: p = 0.8207 | ns |
| | | dTAG13: p = 0.0325 | * |
| GR-P (N) | F(2, 6) = 6.60, p = 0.0305 | KH-103: p = 0.4922 | ns |
| | | dTAG13: p = 0.0684 | # |
| GR-A (N) | F(2, 6) = 20.15, p = 0.0022 | KH-103: p = 0.0120 | * |
| | | dTAG13: p = 0.0820 | # |
| **translational GR isoforms** | **one-way ANOVA** | **Holm-Šídák's: DMSO vs.** | **Summary** |
| GR-α (C) | F(2, 6) = 9.08, p = 0.0153 | KH-103: p = 0.0222 | * |
| | | dTAG13: p = 0.0127 | * |
| GR-α-C3 (C) | F(2, 6) = 46.42, p = 0.0002 | KH-103: p = 0.0002 | *** |
| | | dTAG13: p = 0.0002 | *** |
| GR-α-D3 (C) | F(2, 6) = 101.30, p = < 0.0001 | KH-103: p = < 0.0001 | **** |
| | | dTAG13: p = < 0.0001 | **** |
| **SARS-CoV2-MPro** | **one-way ANOVA** | **Holm-Šídák's: DMSO vs.** | **Summary** |
| Mpro (C) | F(2, 8) = 4.36, p = 0.0524 | KH-103: p = 0.9898 | ns |
| | | dTAG13: p = 0.0825 | # |
| Mpro (N) | F(2, 8) = 16.96, p = 0.0013 | KH-103: p = 0.3505 | ns |
| | | dTAG13: p = 0.0065 | ** |

Supplementary Table. 4_Summary of statistics for GR isoforms and SARS-CoV2-MPro.

| | DEX 18 h | | DEX 2 h | | MIF | | CORT113176 | | KH-103 | |
|---|---|---|---|---|---|---|---|---|---|---|
| gene name | LogFC | FDR | LogFC | FDR | LogFC | FDR | LogFC | FDR | LogFC | FDR |
| CYP2S1 | -1.80 | 0.0000 | -0.12 | 0.3300 | -0.54 | 0.0000 | -0.22 | 0.0160 | -0.11 | 1 |
| CYP4F22 | 5.40 | 0.0000 | 1.50 | 0.0880 | -0.11 | 0.9800 | -0.68 | 0.7700 | 0.42 | 1 |
| CYP26B1 | -0.99 | 0.0000 | -0.52 | 0.0000 | -0.21 | 0.2100 | -0.34 | 0.0032 | -0.20 | 1 |
| CYP51A1 | 0.51 | 0.0000 | 0.02 | 0.9300 | -0.06 | 0.7900 | 0.01 | 0.9700 | 0.04 | 1 |
| CYP24A1 | -0.63 | 0.0000 | -0.74 | 0.0000 | -0.03 | 0.9500 | -0.04 | 0.8400 | -0.01 | 1 |
| CYP1B1 | -0.87 | 0.0000 | -0.39 | 0.0044 | 0.13 | 0.7100 | 2.10 | 0.0000 | 0.00 | 1 |
| CYP4F11 | -1.20 | 0.0000 | -0.09 | 0.8600 | -0.15 | 0.8000 | 0.01 | 0.9900 | -0.13 | 1 |
| CYP1A1 | 1.50 | 0.0000 | 0.85 | 0.0038 | 0.15 | 0.9000 | 4.50 | 0.0000 | 0.17 | 1 |
| CYP26A1 | -1.40 | 0.0000 | -1.30 | 0.0000 | -0.48 | 0.2400 | -0.49 | 0.1600 | -0.56 | 1 |
| CYP3A5 | 0.49 | 0.0000 | 0.35 | 0.0075 | 0.02 | 0.9800 | 0.18 | 0.3900 | 0.21 | 1 |
| CYP4F12 | 0.59 | 0.0000 | 0.43 | 0.0087 | 0.40 | 0.0480 | 0.33 | 0.1100 | 0.28 | 1 |
| CYP27C1 | -0.69 | 0.0009 | -0.42 | 0.1500 | -0.04 | 0.9800 | -0.10 | 0.8700 | -0.23 | 1 |
| CYP2R1 | 0.30 | 0.0062 | -0.01 | 0.9800 | 0.22 | 0.3000 | 0.06 | 0.8700 | 0.08 | 1 |
| CYP4F2 | -0.94 | 0.0270 | 0.42 | 0.5700 | -0.77 | 0.3300 | -0.38 | 0.6700 | -0.35 | 1 |
| CYP4F3 | 0.18 | 0.0430 | 0.40 | 0.0000 | -0.09 | 0.7500 | 0.02 | 0.9500 | 0.93 | 1 |
| CYP20A1 | 0.24 | 0.0470 | 0.11 | 0.7100 | 0.07 | 0.8900 | 0.16 | 0.5000 | 0.94 | 1 |
| not significant | trend # < 0.1 | significant FDR | | | | | | | | |

Supplementary Table. 5_Summary of differential expression analysis for CYPs genes obtained from the RNAseq experiment. All the listed CYPs genes were significantly changed in the DEX 18 h condition. colour labels green: genes selected for boxplot display in Supplementary Fig. 8, red: FDR statistically significant, yellow: FDR # < 0.1, grey: FDR not significant > 0.1).
